# Turnover Regulation of the Rho GTPase Cdc42 by Heat Shock Protein Chaperones and the MAPK Pathway Scaffold Bem4

**DOI:** 10.1101/2021.07.13.452164

**Authors:** Beatriz González, Paul J. Cullen

## Abstract

All cells maintain an axis of polarity that directs the orientation of growth. Cell polarity can be reorganized during development and in response to extrinsic cues to produce new cell types. Rho GTPases are central regulators of cell polarity and signal-dependent cell differentiation. We show here that one of the best understood Rho GTPases, the highly conserved yeast Cdc42p, is turned over by members of the Heat Shock family of Proteins (HSPs). The Hsp40p chaperone, Ydj1p, was required for turnover of Cdc42p by the NEDD4 E3 ubiquitin ligase, Rsp5p, in the proteosome. Cdc42p turnover was regulated by HSPs at high temperatures, and in aging cells where the protein formed aggregates, implicating HSPs in Rho GTPase quality control. We also show that Cdc42p^Q61L^, which mimics the active (GTP-bound) conformation of the protein, was turned over at elevated levels by Ydj1p and Rsp5p. A turnover-defective version of Cdc42p^Q61L^ led to multibudding phenotypes, implicating Cdc42 turnover in singularity in cell polarization. Cdc42p turnover also impacted MAP kinase pathway specificity. A pathway-specific scaffold, Bem4p, stabilized Cdc42p levels, which biased Cdc42p function in one MAPK pathway over another. Turnover regulation of Rho GTPases by HSPs and scaffolds provides new dimensions to the regulation of cell polarity and signal-dependent morphogenesis.

**Significance Statement:** Rho GTPases are switch-like proteins that govern major decisions in cell polarity and signaling in eukaryotes. We elucidate here a pathway that turns over the yeast Rho GTPase Cdc42p, which is mediated by the heat-shock family of proteins (HSPs) and the NEDD4-type E3 ubiquitin ligase Rsp5p. This finding provides a way for HSPs to exert their widespread effects on morphogenetic responses, phenotypic plasticity, and signaling pathways. We also found that turnover of an active version of Cdc42p is critical for modulating cell polarity. Cdc42p turnover also impacted its function in a pathway specific setting, as stabilization of Cdc42p by Bem4p (SmgGDS-type scaffold) influenced the activity of a specific MAPK pathway. HSPs may regulate Rho GTPase turnover in many systems.

## Introduction

Cells establish an axis of polarity to maintain cell shape and to orient cell growth and division. Cell polarity can be reorganized in response to extrinsic cues to impact many biological processes, including cell motility (chemotaxis/chemotropism), development, and differentiation to specific cell types. Ras homology (Rho) GTPases are master regulators of cell polarity and signaling (Etienne-Manneville and Hall, 2002). Rho GTPases cycle between an active GTP-bound conformation that is regulated by guanine nucleotide exchange factors (GEFs), and an inactive GDP-bound conformation this is regulated by GTPase-activating proteins (GAPs). In the GTP-bound conformation, Rho GTPases interact with effector proteins to regulate cellular responses through the reorganization of cytoskeletal elements (Sit and Manser, 2011), the induction of <u>M</u>itogen-<u>A</u>ctivated <u>P</u>rotein <u>K</u>inase (MAPK) pathways and other pathways (Coso et al., 1995; Van Aelst and D’Souza-Schorey, 1997), by interaction with p21-activated (PAK) kinases (Ha and Boggon, 2018; Rane and Minden, 2014; Tetley et al., 2017), and by other mechanisms. Rho GTPases are also regulated by guanine nucleotide dissociation inhibitors [GDIs, (Boulter et al., 2010; Hoffman et al., 2000)] and adaptor proteins (Irazoqui et al., 2003; Lamas et al., 2020) that impact their function, localization, and activity. Rho GTPases can be modified by post-translational modifications (Barthelmes et al., 2020) that include AMPylation (Barthelmes et al., 2020; Woolery et al., 2014), phosphorylation (Chang et al., 2011; Ellerbroek et al., 2003), and ubiquitination (Majolée et al., 2019). Ubiquitin-dependent turnover of Rho GTPases by E3 ubiquitin ligases impacts Rho GTPase function in many settings (Goka and Lippman, 2015; Oberoi-Khanuja and Rajalingam, 2012; Tian et al., 2011; Wang et al., 2003; Wei et al., 2013). Given the multitude of functions regulated by Rho GTPases, an important open question is how Rho GTPases are directed to specific responses in different settings. This question is relevant because mis-regulation of Rho GTPase function is an underlying cause of cancer and other diseases (Haga and Ridley, 2016; Svensmark and Brakebusch, 2019).

<u>H</u>eat <u>S</u>hock <u>P</u>roteins (HSPs) are evolutionarily conserved chaperones that induce protein folding and also promote degradation of proteins that cannot be refolded (Balchin et al., 2016; Whitesell and Lindquist, 2005). HSPs control a broad diversity of morphogenetic responses, including the regulation of cell polarity, morphogenetic plasticity and signal transduction (Calderwood and Gong, 2016; Grad et al., 2011; Pivovarova et al., 2007; Rutherford and Lindquist, 1998; Sun et al., 2021; Walsh et al., 1997; Whitesell and Lindquist, 2005). Despite the fact that HSPs are widely considered to be evolutionary drivers of phenotypic plasticity, the mechanism by which HSPs control cell polarity and signaling – and how they connect to and regulate cell polarity machinery - remains in many cases unclear.

One of the best studied and most well understood Rho GTPases is the highly conserved yeast Cdc42p [yeast and human Cdc42p are 81% identical (Bi and Park, 2012; Kozminski et al., 2000)]. In yeast, Cdc42p is an essential protein and the master regulator of polarity establishment (Bi and Park, 2012; Irazoqui and Lew, 2004; Pringle et al., 1995). Cdc42p also regulates multiple MAPK pathways (Bardwell, 2005; Saito, 2010; Schwartz and Madhani, 2004) that induce different morphogenetic responses through transcriptional regulation of non-overlapping sets of target genes. Several well-defined mechanisms account for specification of shared components, between MAPK pathways, including the utilization of scaffolds (Choi et al., 1994; Marcus et al., 1994; Posas and Saito, 1997; Printen and Sprague, 1994) and degradation of pathway-specific factors (Bao et al., 2004; Chou et al., 2004). However, it remains unclear how Cdc42p and other proteins selectively regulate MAPK pathways that share components. In addition to these roles, Cdc42p is a component of the exocyst complex (Adamo et al., 2001), which controls vesicle delivery to the plasma membrane (Munson and Novick, 2006). Cdc42p also regulates the lysosome/vacuole (Jones et al., 2010; Müller et al., 2001) and has recently been shown to control sealing of the nuclear envelope during cell division (Lu and Drubin, 2020). How a switch-like protein controls all of these biological processes is not clear.

Here, we show that yeast Cdc42p is ubiquitinated and turned over in the proteosome. As this is the first case of Rho GTPase turnover in yeast, we explored the regulation of Cdc42p turnover in this model genetic system and uncovered a role for the NEDD4-type E3 ubiquitin ligase Rsp5p and HSPs in regulating turnover of Cdc42p. HSPs were required for turnover of Cdc42p at high temperatures and under conditions when the protein formed aggregates. Cdc42p aggregates preferentially localized to mother cells, which implicates turnover regulation by HSPs in Rho GTPase quality control. HSPs also mediated turnover of the GTP-bound conformation of the protein, which implicates HSPs in Rho GTPase regulation. By investigating Cdc42p turnover using turnover-defective versions combined with GTP-locked versions of the protein, we identified multi-budding phenotypes, which led to a role for Cdc42p turnover regulation in singularity in cell polarization. We also show that the scaffold Bem4p stabilized Cdc42p levels, which impacted the activity of one Cdc42p-dependent MAPK pathway among several pathways that operate in the same cell type.

## Results

### Cdc42p is Turned Over by Growth at High Temperatures by the E3 Ligase Rsp5p and Hsp40p and Hsp70p Chaperones

Cdc42p is thought to be a stable protein based on immunoblots (IB) analysis, where protein levels have been measured with antibodies to the Cdc42p protein (Adamo et al., 2001; Atkins et al., 2013; Kozminski et al., 2000; Ziman et al., 1993) or antibodies to functional Cdc42p fusion proteins, like green fluorescent protein [GFP (Daniels et al., 2018; Freisinger et al., 2013)]. Consistent with these findings, we found that Cdc42p was stable at 30°C based on treatment with the protein synthesis inhibitor cycloheximide (**Fig. 1A**, CHX). This experiment was carried out in the *Σ*1278b background, which was used for most experiments unless otherwise indicated. Cdc42p turnover at 37°C was not strain-specific and was seen in multiple backgrounds (see below). By comparison, we found that Cdc42p was rapidly degraded in cells incubated at 37°C (**Fig. 1B**). A functional epitope-tagged version of Cdc42p, GFP-Cdc42p (Woods et al., 2016), showed the same pattern of turnover by IB analysis (*Fig. S1, A* and *B*). This result indicates that Cdc42p and GFP-Cdc42p show a similar pattern of turnover, and antibodies to Cdc42p and GFP were used interchangeably in the study. The half-life of Cdc42p was 2.5 min at 37°C and >2h at 30°C (**Fig. 1C**). Therefore, Cdc42p levels change in a condition-specific manner.

**Figure 1.**
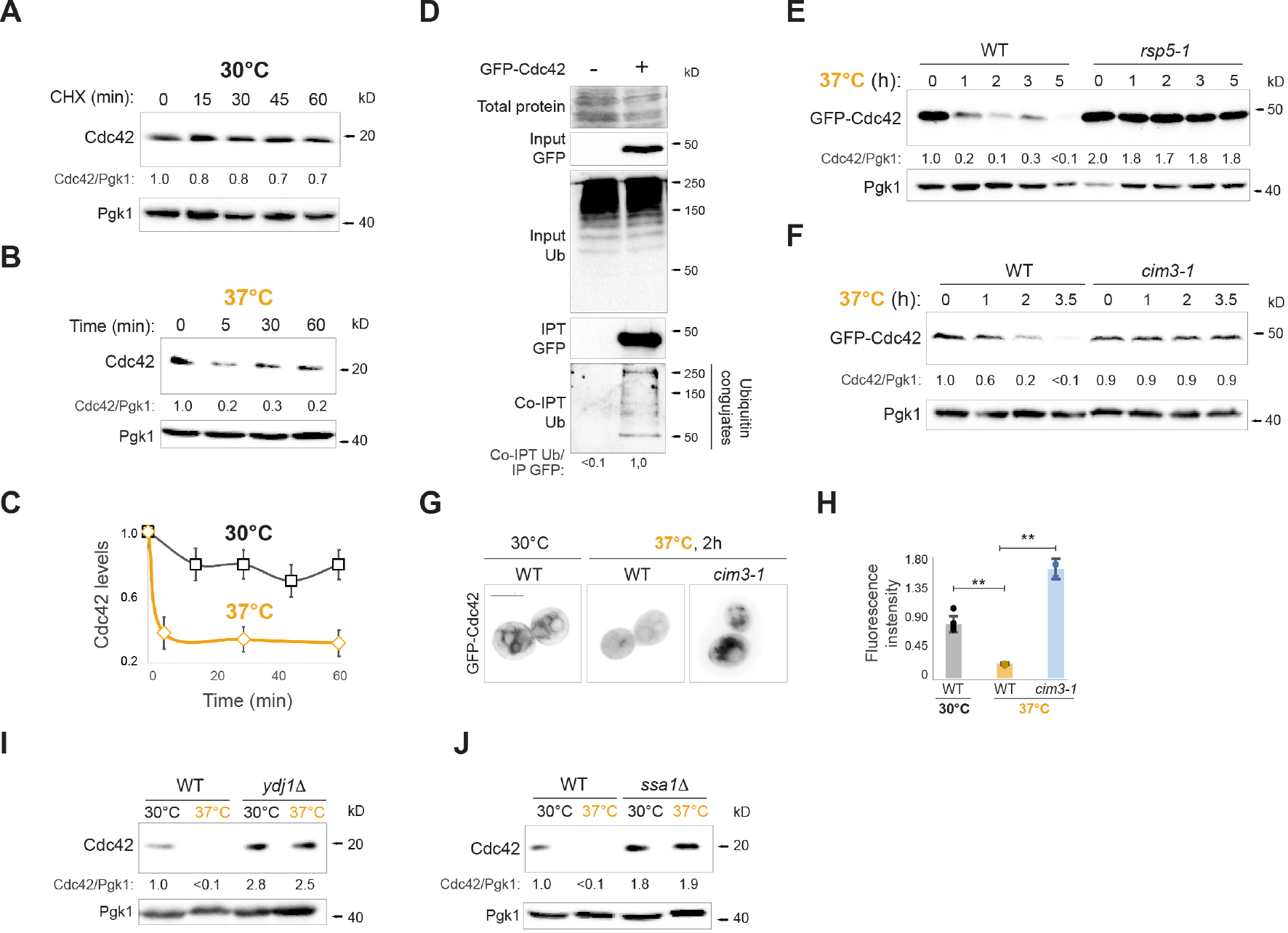
Cdc42p is degraded at 37°C in a Hsp40p-, Hsp70p-, Rsp5p-, and proteasome-dependent manner. **A)** Wild-type (WT) cells (*Σ*1278b background, PC538) were incubated at 30°C in SD+AA media supplemented with 25 µg/ml of CHX and analyzed by IB at the indicated time points. Numbers indicate relative values of Cdc42p compared to Pgk1p at time zero. **B)** The same cells in panel A were incubated at 37°C for the indicated time points and analyzed by IB. **C)** Cdc42p levels relative to Pgk1p at 30°C (black) and at 37°C (yellow). Error bars represent the standard deviation (S.D.) among biological replicates (n=2). **D)** Ubiquitination of Cdc42p was compared in lysates prepared from cells with (+) and without (-) the pGFP-Cdc42p plasmid by Co-IPT. Input and precipitated lysates were blotted against anti-GFP and anti-ubiquitin (Ub) antibodies. Ponceau S staining was used in the Input lysates as a control for protein levels. Numbers indicate relative band intensity of Co-IPT Ub compared to the IPT GFP protein intensity. **E)** Levels of GFP-Cdc42p expressed in WT (S288c background, PC3288) and *rsp5-1* (PC3290) cells grown for 4 h at 30°C and shifted to 37°C. Samples were analyzed at the indicated times. Numbers indicate relative GFP-Cdc42p levels compared to Pgk1p at time zero in WT. **F)** WT (PC5851) and *cim3-1* (PC5852) cells expressing GFP-Cdc42p were grown at 37°C and analyzed by IB at the indicated time points. Numbers indicate the relative levels of GFP-Cdc42p compared to Pgk1p at time zero in WT. **G)** Fluorescence microscopy of WT cells (PC5851) expressing GFP-Cdc42p at 30°C or 37°C and *cim3-1* (PC5852) cells grown at 37°C. Scale bar, 5 µm. **H)** Relative fluorescence intensity of GFP-Cdc42p expressed in cells described in panel G. Error bars represent the standard deviation (S.D.) among biological replicates, n>10, asterisk p-value <0.01. **I)** Cdc42p levels in WT cells (S288c background, PC986) and *ydj1Δ* mutant (PC7556) grown at 30°C for 5 h (30°C) or shifted to 37°C for 2 h (37°C). Numbers indicate relative Cdc42p compared to Pgk1p in WT at 30°C. **J)** Cdc42p levels in WT cells (*Σ*1278b background, PC6016) and *ssa1Δ* mutant (PC7701) grown at 30°C for 5 h (30°C) or shifted to 37°C for 2 h (37°C). Numbers indicate relative levels of Cdc42p compared to Pgk1p in WT at 30°C.

Many proteins, including monomeric and heterotrimeric GTPases (Dohlman and Campbell, 2019), are turned over by ubiquitin modification; however, Rho GTPase turnover has not been explored in yeast. Immunoprecipation (IPT) of GFP-Cdc42p showed cross-reactivity to anti-ubiquitin antibodies, indicating that the protein is ubiquitinated (**Fig. 1D**). E3 ubiquitin ligases control protein turnover by the covalent attachment of ubiquitin onto substrates (Buetow and Huang, 2016). A candidate approach identified Rsp5p, a member of the NEDD4 (<u>N</u>europrecursor cell <u>E</u>xpressed <u>D</u>evelopmentally <u>D</u>ownregulated 4) family of E3 ubiquitin ligases (Ingham et al., 2004), as being required for Cdc42p turnover. Rsp5p is an essential protein in yeast (Huibregtse et al., 1995). A temperature-sensitive allele, *rsp5-1*, with a point mutation in the catalytic <u>H</u>omologous to <u>E</u>6AP <u>C-T</u>erminus (HECT) domain (L733S) (Fisk and Yaffe, 1999; Wang et al., 1999), was defective for GFP-Cdc42p turnover at 37°C (**Fig. 1E**, S288c background). This result was confirmed using antibodies to the Cdc42p protein (*Fig. S1C*). Rsp5p contains three WW domains that recognize PY (PPxY) motifs on substrate proteins (Gajewska et al., 2001). A version of Rsp5p lacking a functional WW3 domain, Rsp5p^W451G^, showed stabilization of Cdc42p (*Fig. S1D*). Rsp5p regulates the turnover of cytosolic proteins in the proteasome (Brückner et al., 2011; Challa et al., 2021; Kowalski et al., 2018) and integral-membrane proteins by vesicular trafficking to the lysosome/vacuole (Katzmann et al., 2001; Lin et al., 2008). The proteosome was evaluated using the temperature-sensitive *cim3-1* mutant (Ghislain et al., 1993). The *cim3-1* mutant showed elevated levels of Cdc42p (**Fig. 1F**), which indicates that the proteasome is required for Cdc42p turnover. As an independent test of Cdc42p protein levels, GFP-Cdc42p levels were measured by fluorescence microscopy. GFP-Cdc42p can be found at sites of growth and internal compartments (Richman et al., 2002). GFP-Cdc42p showed reduced levels at 37°C (**Fig. 1**, **G** and **H**), and the levels of the protein accumulated in the *cim3-1* mutant (**Fig. 1**, **G** and **H**). Therefore, Cdc42p is turned over by Rsp5p in the proteasome.

Rsp5p recognizes PY motifs on target proteins through its WW domains. Cdc42p does not contain a PY motif, suggesting that an adaptor might mediate its recognition. It has been shown that heat shock proteins (HSPs) can function as adaptor proteins that mediate the turnover of proteins lacking PY motifs in an Rsp5p-dependent manner (Fang et al., 2014). One HSP adaptor for Rsp5p is the Hsp40p chaperone, Ydj1p (Fang et al., 2014). Ydj1p was required for turnover of Cdc42p at 37°C (**Fig. 1I**). Hsp40p chaperones function as co-chaperones for Hsp70p chaperones (Glover and Lindquist, 1998; Qiu et al., 2006). Ydj1p functions as a co-chaperone for Hsp70 proteins (Becker et al., 1996), which also regulate protein degradation by the ubiquitin-proteasome system (Guerriero et al., 2013; Lee do et al., 2016). Cells lacking one Hsp70p member, Ssa1p, were defective for Cdc42p turnover (**Fig. 1J**), indicating that, like Ydj1p, Ssa1p is required for turnover of Cdc42p at 37°C. The protein kinase C (PKC) pathway, which regulates the response to temperature stress (Heinisch and Rodicio, 2018; Kamada et al., 1995), was not required for turnover of Cdc42p at 37°C (*Fig. S1F*). Therefore, Cdc42p is turned over by a mechanism that involves Hsp40p and Hsp70p chaperones, the E3 ligase Rsp5p, and the proteasome.

### GTP-Locked Version of Cdc42p is Ubiquitinated and Turned Over by the NEDD4 E3 Ubiquitin Ligase Rsp5p in the Proteosome

Cdc42p turnover also occurred to some degree at 30°C, which required Ydj1p (**Fig. 1I**), Ssa1p (**Fig. 1J**), Rsp5p (**Fig. 1E**, time zero and *Fig. S1C*), its WW3 domain (*Fig. S1D*), and the proteasome, based on treatment of cells with the proteosome inhibitor MG132 (Fenteany et al., 1995; Finley et al., 2012) (*Fig. S1F*). This prompted us to investigate whether the turnover of Cdc42p impacts its biological function or activity. To address this possibility, turnover of GFP-Cdc42p was compared to GFP-Cdc42p^Q61L^, which mimics the GTP-bound conformation (Ziman et al., 1991). Theproteins were expressed from the *CDC42* promoter from plasmids in strains containing a chromosomal copy of *CDC42*. This allowed examination of turnover of active Cdc42p in viable cells, because cells expressing Cdc42p^Q61L^ as the sole copy are lethal (Ziman et al., 1991). By IB analysis, GFP-Cdc42p^Q61L^ was less stable than GFP-Cdc42p (**Fig. 2A**, Cdc42 and **Fig. 2B**, Q61L). GFP-Cdc42p^Q61L^ had a half life of <30 min (**Fig. 2C**). Likewise, strains containing plasmids expressing GFP-Cdc42p^Q61L^ (*Fig. S2A*) and GFP-Cdc42p^G12V^ (*Fig. S2B*), which mimic the GTP-bound conformation of the protein, showed lower steady-state levels of Cdc42p. Expression of these proteins was not toxic in the S288c or *Σ*1278b backgrounds, which differs from previous reports (Woods et al., 2016; Ziman et al., 1991), perhaps because the GFP fusion reduces toxicity. The degradation profile of GFP-Cdc42p^Q61L^ appeared biphasic rather than exponential (**Fig. 2B**), which might reflect multiple mechanisms underlying turnover. By fluorescence microscopy, cells expressing GFP-Cdc42p were brighter than cells expressing GFP-Cdc42p^Q61L^ (**Fig. 2D**). The difference in Cdc42p levels by fluorescence intensity (**Fig. 2E**, 7-fold) was similar to the difference in band intensity seen by IB analysis (*Fig. S2A,* 10-fold). Cells expressing GFP-Cdc42p^Q61L^ accumulated high molecular weight products that were suggestive of ubiquitin conjugates (**Fig. 2B**, upper right panel). Co-IPT analysis showed that GFP-Cdc42p^Q61L^ was ubiquitinated at 40-fold higher levels than GFP-Cdc42p (**Fig. 2F**). Therefore, a version of Cdc42p that mimics the active GTP-bound conformation is highly ubiquitinated and turned over in yeast.

**Figure 2.**
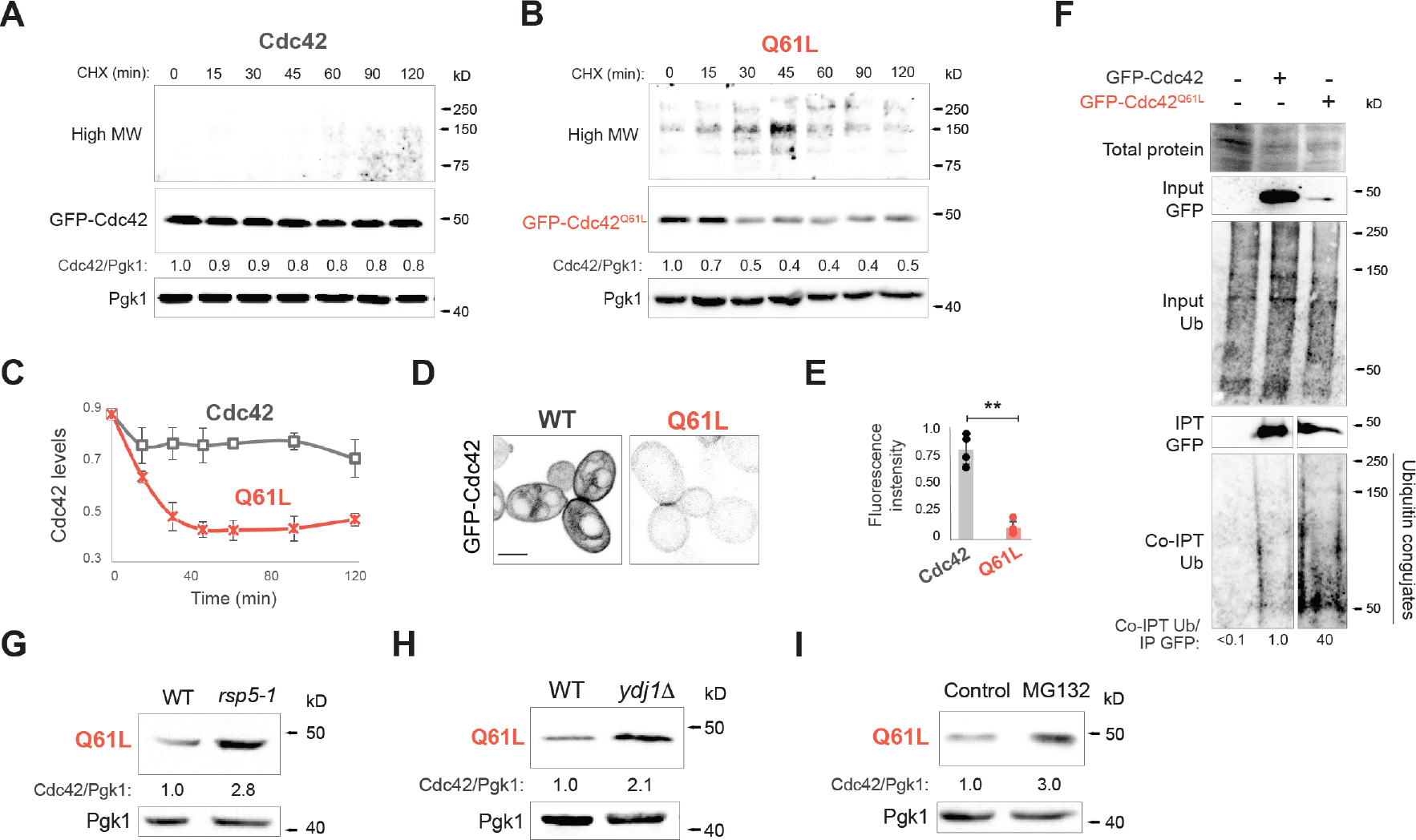
GTP-locked Cdc42p is rapidly degraded in a Ydj1p-, Rsp5p- and proteasome-dependent manner. **A)** WT (PC538) cells expressing a plasmid containing GFP-Cdc42p (Cdc42, PC6454) were incubated in SD-URA media containing 25 µg/ml of CHX and analyzed by IB at the indicated time points. Numbers indicate the relative band intensity to GFP-Cdc42p levels at time 0. Top panel corresponds to longer exposure time of anti-GFP blot. **B)** Same as panel A except cells contain plasmid GFP-Cdc42p^Q61L^ (Q61L, PC7458). **C)** Levels of GFP-Cdc42p (Cdc42, black, set to 1 for time zero) and GFP-Cdc42p^Q61L^ (Q61L, orange, also set to 1 for time zero). Levels of GFP-Cdc42p^Q61L^ at time zero is 5-fold lower than GFP-Cdc42p. Error bars represent the standard deviation (S.D.) among biological replicates (n=2). **D)** Fluorescence microscopy of WT cells expressing GFP-Cdc42p (WT) or GFP-Cdc42p^Q61L^ (Q61L). Scale bar, 5 µm. **E)** Relative fluorescence intensity of cells expressing GFP-Cdc42p (Cdc42) or GFP-Cdc42p^Q61L^ (Q61L). Error bars represent the standard deviation (S.D.) among biological replicates, n>25, asterisk p-value <0.01. **F)** Ubiquitin levels of GFP-Cdc42p and GFP-Cdc42p^Q61L^ by Co-IPT. Input and precipitated lysates were blotted against anti-GFP and anti-ubiquitin antibodies. Ponceau S staining indicates control for protein levels. IPT GFP blot for GFP-Cdc42p^Q61L^ corresponds to a longer exposure time to equal the amount of precipitated protein to GFP-Cdc42p. Same exposure time was applied and is showed in the Co-IPT blot for GFP-Cdc42p^Q61L^. Numbers indicate relative band intensity of Co-IPT Ub compared to IPT GFP intensity. **G)** GFP-Cdc42p^Q61L^ levels in WT (PC3288) and *rsp5-1* (PC3290) cells at the permissive temperature, 30°C. Numbers indicate relative levels of GFP-Cdc42p. **H)** GFP-Cdc42p^Q61L^ levels in WT (S288c background, PC986) cells and *ydj1Δ* (PC7657). Numbers indicate the relative levels of GFP-Cdc42p compared to Pgk1p at time zero. **I)** WT cells (PC538) containing GFP-Cdc42p^Q61L^ supplemented with 0.5% ethanol (control) or 75μM MG132 were incubated for 2 h. Numbers indicate relative levels of GFP-Cdc42p compared to Pgk1p.

We analyzed whether the proteins involved in turnover at 37°C also regulate the degradation of Cdc42p^Q61L^. Like at 37°C, the *rsp5-1* mutant was defective for turnover of GFP-Cdc42p^Q61L^ (**Fig. 2G**). Rsp5p^W451G^, which lacks a functional WW3 domain, was also defective for turnover of GFP-Cdc42p^Q61L^ (*Fig. S2C*). The turnover of GFP-Cdc42p^Q61L^ also required Ydj1p (**Fig. 2H**, 2-fold) indicating that HSPs regulate the turnover of active Cdc42p. GFP-Cdc42p^Q61L^ was stabilized by addition of MG132 to cells (**Fig. 2I**, 3-fold) but not in cells lacking the vacuolar protease Pep4p (*Fig. S2D*). In fact, cells lacking Pep4p showed reduced levels of Cdc42p^Q61L^, indicating that vacuolar degradation of proteins might negatively impact Cdc42p degradation by the proteasome. Therefore, incubation of cells at 37°C and Cdc42p^Q61L^ are turned over by a mechanism that involves Hsp40p, Rsp5p, and the proteosome.

**Figure 3.**
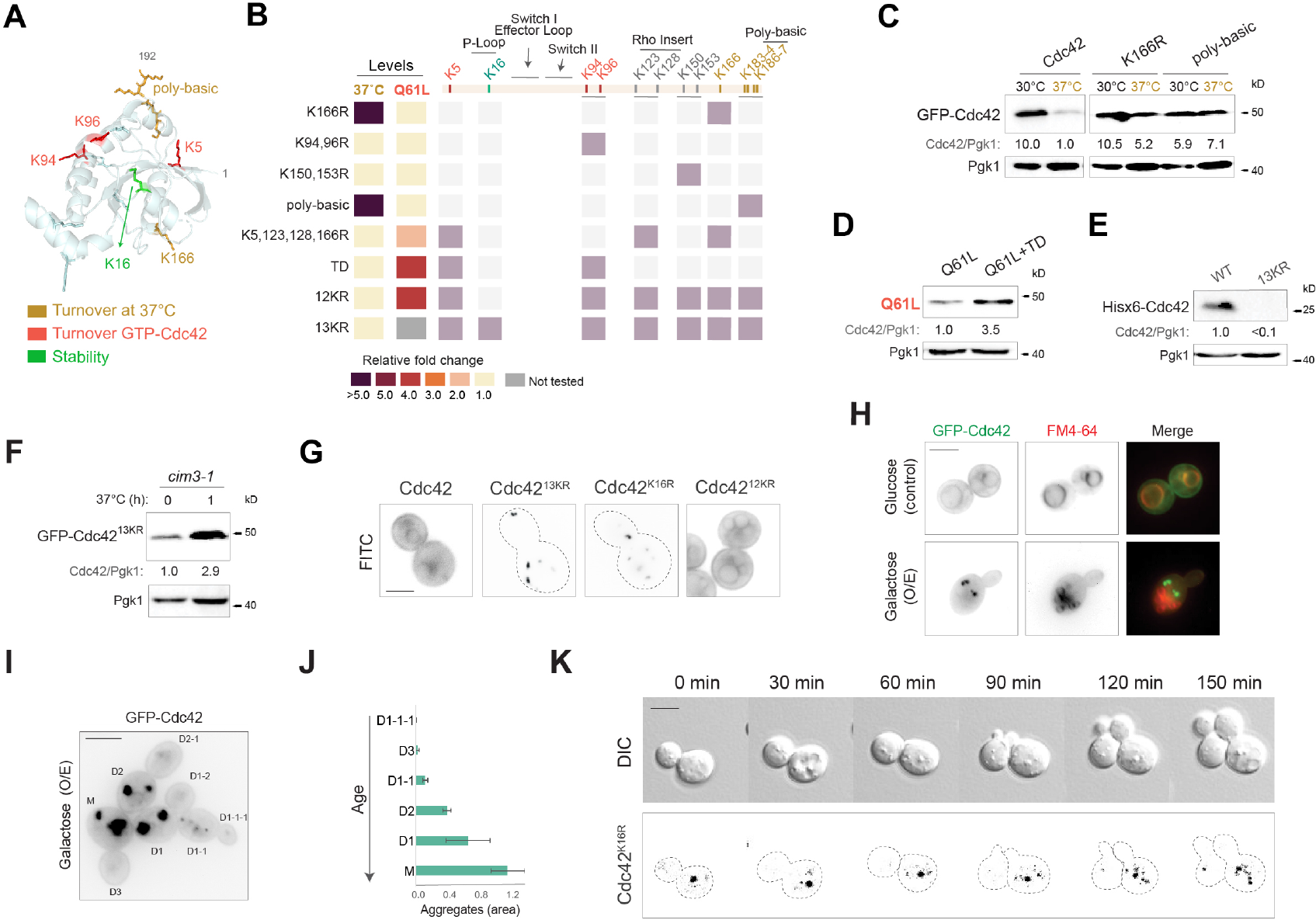
Lysine residues are required for Cdc42p turnover and stability. Alterations of Cdc42p lead to the formation of aggregates in mother cells. **A)** The yeast Cdc42p protein sequence was overlaid onto the crystal structure of human Cdc42p using the Expasy web server SWISS-MODEL (https://swissmodel.expasy.org) (Nassar et al., 1998). Yellow refers to lysines involved in Cdc42p turnover at 37°C, red refers to lysines involved GTP-Cdc42p turnover and green indicates lysine involved in protein stability. **B)** Diagram of the Cdc42p protein which shows the location of 13 lysine residues. Heat map indicates lysines substitutions to arginines in Cdc42p. Colors of alleles are described in panel 3A. Color code represents the fold change of Cdc42p-alleles amount compared to wild-type Cdc42p. TD, turnover deficient, K5,94,96R. **C)** GFP-Cdc42p levels of wild-type (WT) cells expressing GFP-Cdc42p, GFP-Cdc42p^K166R^, or GFP-Cdc42p^K183,184,186,187R^ (polybasic) grown and at 30°C for 5 h (30°C) or shifted to 37°C for 1 h (37°C). Anti-GFP and anti-Pgk1p antibodies were used. Numbers indicate the relative levels of GFP-Cdc42p compared to Pgk1p. Here, Cdc42p levels at 37°C were set to a value of 1. **D)** GFP-Cdc42p levels of WT cells expressing GFP-Cdc42p^Q61L^ or GFP-Cdc42p^Q61L+TD^. Anti-GFP and anti-Pgk1p antibodies were used. Numbers indicate the relative levels of GFP-Cdc42p compared to Pgk1p. **E)** WT cells expressing His6x-Cdc42p or His6x-Cdc42p^13KR^ (His6x-Cdc42p^K5,16,94,96,123,128,150,153,166,183,184,186,187R^). Anti-Cdc42p and anti-Pgk1p antibodies were used. Numbers indicate the relative levels of GFP-Cdc42p compared to Pgk1p. F) *cim3-1* cells _expressing GFP-Cdc42p or GFP-Cdc42p_13KR _(GFP-Cdc42p_K5,16,94,96,123,128,150,153,166,183,184,186,187R_)._ Anti-GFP and anti-Pgk1p antibodies were used. Numbers indicate the relative levels of GFP-Cdc42p compared to Pgk1p**. G)** Fluorescence microscopy of *cim3-1* cells expressing GFP-Cdc42p, GFP-Cdc42p^13KR^, GFP-Cdc42p^12KR^ or GFP-Cdc42p^K16R^ grown at 30°C for 5 h and shifted to 37°C for 1 h. Scale bar, 5 µm. **H)** Fluorescence microscopy of WT cells expressing GFP-Cdc42p (control) or pPGAL1-GFP-linker-*CDC42P* (O/E, overexpression) grown for 6 h in YPED or YPE-GAL, respectively and stained with the lipophilic FM4-64 dye. Scale bar, 5 µm. **I)** Fluorescence microscopy of WT cells expressing pPGAL1-GFP-linker-*CDC42P* (O/E, overexpression) grown for 8 h in YPE-GAL. M refers to mother cell and D to daughter cell. Numbers indicate cell linage. Scale bar, 5 µm. **J)** Relative quantification of the aggregates area from cells of panel 3I and S3I. **K)** Time-lapse microscopy of *cim3-1* cells expressing GFP-Cdc42p^K16R^ grown at 36°C for 150 min. Scale bar, 5 µm.

### Analysis of the Role of Lysines in Cdc42p Turnover and Stability

We next investigated the role that lysine residues play in regulating Cdc42p turnover. Lysine residues typically serve as sites for ubiquitination (Chau, 1989). Cdc42p has 13 lysines, twelve of which are surface exposed (**Fig. 3A**, Movie 1). Site-directed mutagenesis was used to change lysines in Cdc42p to arginines in groups (**Fig. 3B**), because non-preferred lysines can be used when the preferred lysine is absent. Several lysine substitutions rescued the reduced levels of Cdc42p seen at 37°C among a larger panel of combinations tested (*Fig S3A*). These proteins were expressed as GFP-fusions from plasmids in otherwise wild-type cells. Specifically, Cdc42p^K166R^, and Cdc42p^K183R, K184R, K186R, K187R^, which contains lysine substitutions in the poly-basic domain, resulted in higher levels of Cdc42p at 37°C (**Fig. 3**, **B** and **C**). Interestingly, K166 is a site of ubiquitination for Rac1 (Zhao et al., 2013) and Cdc42p (Murali et al., 2017) in humans. Site-directed mutagenesis was also used to change lysines in GFP-Cdc42p^Q61L^. Here, a different combination of lysine substitutions was required for turnover (**Fig. 3, B** and **D**, Cdc42p^Q61L, K5R, K94R, K96R^ or Cdc42p^Q61L+TD^ for <u>T</u>urnover <u>D</u>efective) among a larger set of combinations tested (*Fig S3B*). Cdc42p ^K5R, K94R, K96R^ did not affect the localization of Cdc42p and would not be expected to affect the activity of the protein. We note that K6 (K5 in yeast) is a site of ubiquitination for RhoA (Ozdamar et al., 2005). Therefore, different lysine residues mediate the turnover of Cdc42p at 37°C and the GTP-bound species of the protein.

A version of Cdc42p lacking all 13 lysines was also constructed (Hisx6-Cdc42p^13KR^, the Hisx6 epitope also lacks lysine residues). Unexpectedly, Hisx6-Cdc42p^13KR^ was found at reduced levels (**Fig. 3E**), which indicates that Cdc42p lacking all lysine residues is not stable. Here, instability of Cdc42p superseded the requirement for lysines for turnover at 37°C and Cdc42p^Q61L^. Moreover, this result indicates that Cdc42p can be turned over in a lysine-independent manner, which for some proteins involves other amino acid residues (Ben-Saadon et al., 2004; McDowell and Philpott, 2013; Tait et al., 2007). GFP-Cdc42p^13KR^ accumulated in the *cim3-1* mutant (**Fig. 3F**), indicating that this version of the protein is degraded in the proteosome. The stability defect was traced to a single lysine (K16) that is not surface exposed and that has previously be shown to be required for cell viability (Kozminski et al., 2000) and is presumably required for Cdc42p to be functional. Combinations of K16R with other lysine substitutions destabilized Cdc42p by IB analysis (*Fig. S3C*). We also generated Cdc42p^12KR^, which contains a single lysine at K16. Cdc42p^12KR^ was present at normal levels in the cell (*Fig. S3A*), confirming that K16 is the key residue that is critical for stabilization of the protein. Therefore, the stability of Cdc42p is impacted by lysines that promote turnover, and one lysine (K16), which stabilizes the protein.

### Cdc42p Forms Aggregates When Overproduced or Mis-folded That Are Asymmetrically Localized to Mother Cells

Many cytosolic proteins and mRNA-protein complexes can form aggregates in the cell, which can occur as a result of cellular stresses and aging (Cabrera et al., 2020; Hill et al., 2017; Khong and Parker, 2020; Samant et al., 2018). The aggregation of proteins can be induced by misfolding, which is a major cause of folding-related neurodegenerative disorders in humans (Balchin et al., 2016; Dobson, 2002). The accumulation of GFP-Cdc42p^13KR^ (expressed from a plasmid in cells containing a genomic copy of *CDC42*) in the *cim3-1* mutant led to aggregation of the protein (**Fig. 3G**). This result supports the idea that this version of the protein is prone to misfolding, and its degradation by the proteasome prevents aggregation. GFP-Cdc42p^K16R^ also formed aggregates (**Fig. 3G**), and induced aggregate formation in combination with other lysine substitutions (*Fig. S3, D* and *E*). As discussed above, these changes are expected to led to nonfunctional versions of the protein. Cdc42p^12KR^, which does not contain the K16R mutation, did not form aggregates under this condition (**Fig. 3G**).

We next asked whether the wild-type Cdc42p protein can form aggregates. Growth at 37°C did not induce aggregation of Cdc42p, but overexpression of the *GFP-CDC42* gene from an inducible promoter (P*GAL1*) induced aggregate formation (**Fig. 3H**). Similarly, GFP-Cdc42p formed aggregates in cells lacking Ydj1p, where the protein accumulates at early time points (*Fig. S3F*). Aggregates varied in number and size and showed a cytosolic pattern that was distinct from the lipophilic dye FM4-64 (**Fig. 2H**). Aggregates of misfolded proteins can be kept in larger inclusions in certain locations within the cell to prevent harmful interactions and facilitate degradation. Under conditions that promoted aggregate formation, Cdc42p showed elevated turnover, as determined by shifting cells to media containing glucose to induce promoter shut off (*Fig. S3*, *G* and *H*). Aggregates precursors are also retained in aging mother cells, by a HSP-dependent mechanism (Saarikangas et al., 2017), to ensure that properly folded proteins are present in daughter cells to promote cellular rejuvenation (Hill et al., 2017). When overexpressed, Cdc42p aggregates preferentially accumulated in older cells (**Fig. 3**, **I** and **J**, *Fig. S3I*), Here, the filamentous strain background (*Σ*1278b) facilitated determination of cell lineage, because adhesive cells fail to separate. Similarly, GFP-Cdc42p^K16R^ was enriched in mother cells throughout multiple cell divisions (Movie 2, Cdc42p; Movie 3, Cdc42p^K16R^; **Fig. 3K**). Therefore, the Cdc42p protein can form aggregates when overproduced, by overexpression or if turnover is impaired, and if the protein is damaged in a way that compromises stability. The retention of aggregated Cdc42p in aging cells in these settings may provide a mechanism for the rejuvenation of cell polarity.

### Turnover of GTP-Cdc42p Is Required for Singularity in Cell Polarization

To further analyze the biological significance Cdc42p turnover, versions of GTP-Cdc42p that were defective for turnover were examined. Cells expressing a wild-type version of Cdc42p that cannot be turned over (Cdc42p^TD^) as the only copy of *CDC42* in the cell were not viable (**Fig. 4A**) indicating that the turnover of Cdc42p might have an essential function. Alternatively, Cdc42p^TD^ might not be functional; however, when combined with Cdc42p^Q61L^, gain-of-function phenotypes are seen in cell polarity and MAPK signaling (see below), which suggests that these changes do not compromise the function of the protein. In line with the idea that turnover of Cdc42p is critical for function, the *rsp5-1* (*Fig. S4A*) and *ydj1Δ* (*Fig. S4B*) mutants harboring GFP-Cdc42p^Q61L^ exhibited growth defects, presumably due to the accumulation of GTP-bound Cdc42p, which would be expected to compromise viability (Ziman et al., 1991). The terminal phenotype of cells expressing Cdc42p^TD^ showed abnormal cell morphologies by microscopic examination, including large cells with multiple buds (**Fig. 4A**, bottom panels). The *rsp5-1* (*Fig. S4C*) and *ydj1Δ* (*Fig.S4E*) mutants harboring Cdc42p^Q61L^ also exhibited abnormal morphologies, including the formation of large cells (*Fig. S4C* and *D*). The *rsp5-1* and *ydj1Δ* mutants showed different types of cell morphologies, which may occur because these proteins may have non-overlapping client proteins and targets (Meacham et al., 1999). Importantly, turnover of a GTP-locked allele of Cdc42p may be critical for the overall regulation of cell polarity.

**Figure 4.**
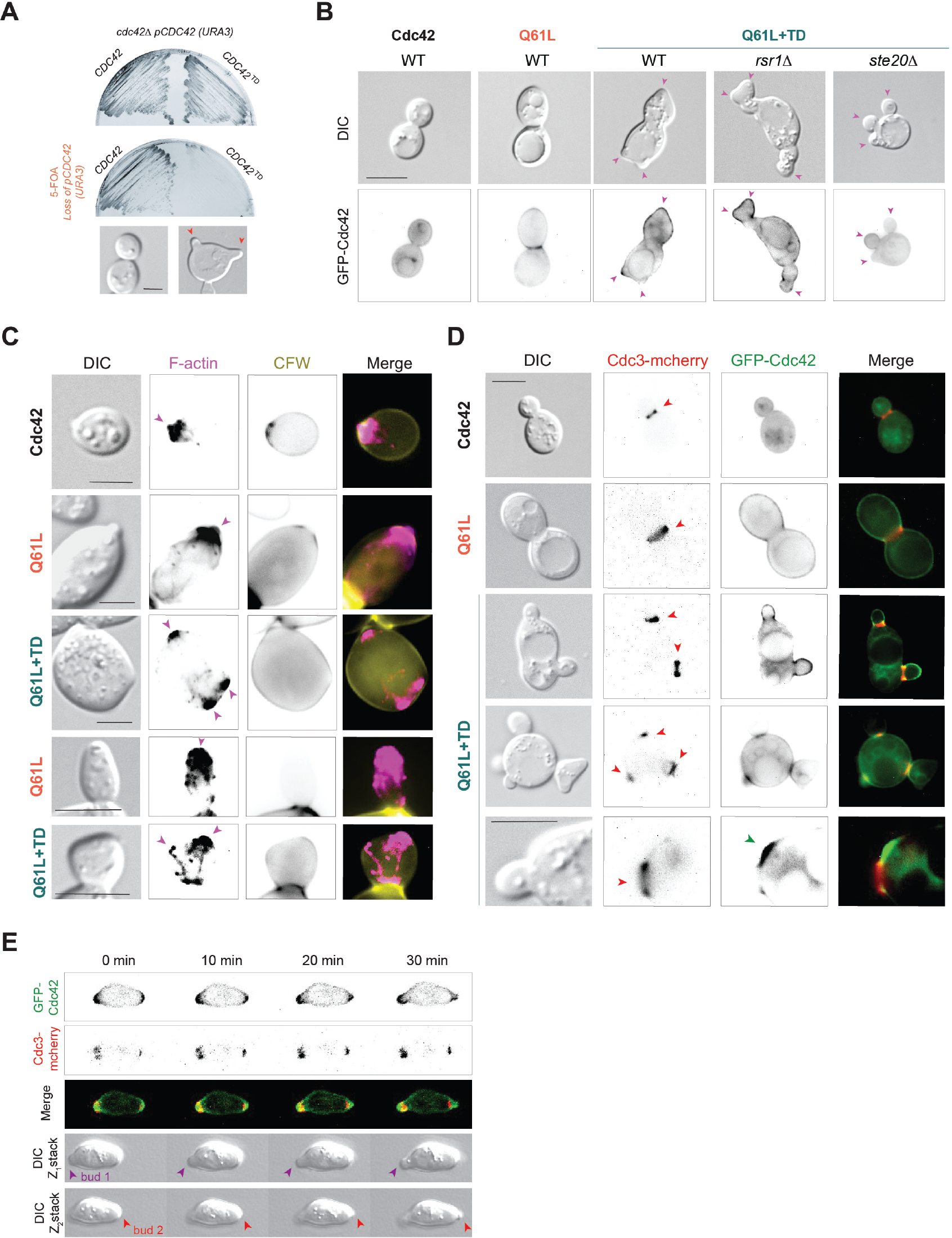
Role of Cdc42p turnover in viability and singularity in polarization. **A)** Top, *cdc42Δ* strain harboring the pRS316-GFP-Cdc42 (*URA3*) and pRS315-GFP-Cdc42 (*LEU2*) or pRS315-GFP-Cdc42^TD^ (TD, K5,94,96R; *LEU2*) were examined for growth on SD-URA-LEU and 5-FOA media. Bottom, morphology of *cdc42Δ* strain harboring the pRS316-GFP-Cdc42 (*URA3*) and pRS315-GFP-Cdc42 (*LEU2*) or pRS315-GFP-Cdc42^TD^ (TD, K5,94,96R; *LEU2*) after 2 days on 5-FOA media. Red arrows indicate multiple buds. Scale bar, 5 µm. **B)** WT cells expressing GFP-Cdc42p (Cdc42), GFP-Cdc42p^Q61L^ (Q61L) or GFP-Cdc42p^Q61L+K5,94,96R^ (Q61L+TD) and *rsr1Δ* and *ste20Δ* mutants expressing GFP-Cdc42p^Q61L+K5,94,96R^ (Q61L+TD). Red arrows indicate multiple growth projections. Scale bar, 5 µm. **C)** WT cells expressing GFP-Cdc42p (Cdc42), GFP-Cdc42p^Q61L^ (Q61L) or GFP-Cdc42p^Q61L+K5,94,96R^ (Q61L+TD) were grown at 30°C for 5 h and fixed with 4% formaldehyde. Cells were stained with Phalloidin-Atto 532 (F-actin, magenta) and fluorescent brightener #28 (CFW, calcofluor white, yellow, cell wall). Magenta arrows refer to multiple actin polymerization. Scale bar, 5 µm. **D)** Fluorescence microscopy of Cdc3-mcherry cells expressing GFP-Cdc42p (Cdc42), GFP-Cdc42p^Q61L^ (Q61L) or GFP-Cdc42p^Q61L+K5,94,96R^ (Q61L+TD). Cdc3-mcherry (red) is a septin ring marker. Red arrows refer to multiple septin rings. Scale bar, 5 µm. **E)** Confocal time-lapse microscopy of WT cells expressing GFP-Cdc42p^Q61L+K5,94,96R^ (Q61L+TD). Cdc3-mcherry (red) is a septin ring marker. Different color arrows refer to different buds. Scale bar, 5 µm

Introduction of Cdc42p^Q61L+TD^ in wild-type cells that contain a copy of the *CDC42* gene was not toxic and did not cause lethality, which allowed evaluation of the role of Cdc42p turnover on cell polarity regulation. Cells expressing Cdc42p^Q61L+TD^ induced the formation of misshapen cells and cells with multiple growth projections (**Fig. 4B**, *Fig. S4F,* Cdc42p^TD^ alone has modest phenotypes). In the GTP-bound state, Cdc42p interacts with the formin, Bni1p (Evangelista et al., 1997), and two p21-activated kinases (PAKs), Ste20p and Cla4p (Evangelista et al., 2000; Lechler et al., 2001), which collectively direct actin cable assembly at nascent sites. Immunofluorescence staining showed that F-actin was polarized at multiple sites [**Fig. 4C**, arrows; GFP-Cdc42p, <0.1% of cells showed actin at multiple sites (n= 250), GFP-Cdc42p^Q61L^, 0.9% (n= 152), GFP-Cdc42p^Q61L+TD^, 15.8% (n= 184)]. Cdc42p also interacts with Gic1p and Gic2p (Brown et al., 1997; Iwase et al., 2006), which control assembly of the septin ring that creates a barrier to restrict growth to the emerging bud (Daniels et al., 2018; Okada et al., 2013). Cells expressing Cdc42p^Q61L+TD^ had multiple septin rings and formed multiple buds [**Fig. 4D**; GFP-Cdc42p, <0.1% of cells formed multiple buds (n= 229), GFP-Cdc42p^Q61L^, 1.7% (n= 114), Cdc42p^Q61L+TD^ 16.5% (n= 164); *Fig. S4G*]. Time-lapse microscopy showed that in some cells containing two sets of septin rings, buds grew at the same time (**Fig. 4E,** 24%), while in other cells, buds grew in sequence (Movie 4, Cdc42p; Movie 5, Cdc42p^Q61L^; Movie 6, Cdc42p^Q61l+TD^, 76%). New growth sites initiated by GFP-Cdc42p^Q61L+TD^ occurred outside the septin ring (**Fig. 4D**, bottom panel). Multiple projections also formed inside buds (Movie 7). Here, actin cables extended to multiple sites (**Fig. 4C**, bottom two panels), which led to the formation of multiple projections within the bud.

Cells expressing Cdc42p^Q61l+TD^ also exhibited lysis (e.g. Movie 7, arrow). Cell lysis might account for the growth defect of these cells (**Fig.** 4A, *Fig. S4A* and *B*), which is known to occur in cells expressing high levels of Cdc42p^Q61L^ (Ziman et al., 1991) or Cdc42p^G12V^ (Gulli et al., 2000). It has previously been suggested that GTP-locked versions of Cdc42p cause ‘wounds’ in the cell wall that result in the formation of multiple growth sites (Woods and Lew, 2019). In addition, Cdc42p^Q61L^ induced an increase in cell size. Multibudded cells can arise in large cells as a result of altered dosage of polarity proteins (Chiou et al., 2021); however, Cdc42p^Q61l+TD^ cells were same size as Cdc42p^Q61L^ cells, which indicates that this is not the case. Thus, failure to turn over a GTP-locked version of Cdc42p results in the formation of multiple sites of growth.

Cdc42p is required for cell polarization (Wedlich-Soldner et al., 2003) through a well-understood series of positive and negative feedback loops (Caviston et al., 2002; Irazoqui et al., 2003; Slaughter et al., 2009; Woods et al., 2016). Multi-budding is indicative of a defect in singularity in polarization, which ensures that polarity establishment occurs once per cell cycle (Caviston et al., 2002; Goryachev and Leda, 2017; Irazoqui et al., 2003; Slaughter et al., 2009; Woods et al., 2016). The turnover of GTP-bound Cdc42p may help to resolve competition at sites where cell polarization occurs and provide negative feedback to ensure that only a single bud is formed at a time. In order to produce a new cell, Cdc42 is activated at bud sites by spatial cues that mark the cell surface at the end of G1 (Chant and Pringle, 1995; Howell and Lew, 2012; Miller et al., 2020; Moran et al., 2019). When spatial signals are disrupted, Cdc42p can induce cell polarization by a process known as symmetry breaking (Irazoqui et al., 2003; Kozubowski et al., 2008; Martin, 2015; Woods et al., 2016). During this process, Cdc42p is activated at random sites and becomes concentrated at a single site by a positive feedback. In cells lacking the ability to grow at bud sites, [*rsr1Δ* (Park et al., 2002)], GFP-Cdc42p^Q61L+TD^ induced more growth sites than wild type, indicating that bud sites function to restrict growth from the accumulation of GTP-bound Cdc42p (**Fig. 4B**). A downstream effector of Cdc42p, Bni1p, was required for polarization in buds but not for multi-budding (data not shown), which fits with the idea that Bni1p promotes actin polymerization but is not required for bud emergence (Woods et al., 2016). Another effector of Cdc42p, the PAK Ste20p, was not required for multi-budding phenotypes (**Fig. 4B**). Therefore, the accumulation of active Cdc42p induces growth preferentially at bud sites by the utilization of a subset of effector proteins.

### Stabilization of Cdc42p by Bem4p Results in Activation of a Specific MAPK Pathway

In addition to regulating bud emergence, Cdc42p also regulates MAPK pathways. These pathways can share components (**Fig. 5A**). One pathway mediates the response to mating pheromone (Simon et al., 1995), in which complementary cell types mate to form diploids (Alvaro and Thorner, 2016; Bardwell, 2005). Another pathway regulates filamentous growth (Peter et al., 1996; Truckses et al., 2006; Wu et al., 1998), a cell differentiation response that occurs in many fungal species including pathogens (Fisher et al., 2020; Mitchell, 1998), which in yeast occurs in response to limiting nutrients (Gimeno et al., 1992; Jin et al., 2008; Lorenz and Heitman, 1997). A third Cdc42p-dependent MAP kinase pathway regulates the response to osmotic stress (Hohmann, 2015; Saito and Posas, 2012; Tatebayashi et al., 2006). Attempts to evaluate the roles of Rsp5p and Ydj1p in regulating MAPK pathway signaling showed pleotropic phenotypes, perhaps because filamentous growth and the fMAPK pathway are controlled by many proteins and pathways in an integrated network (Chavel et al., 2014), and because HSPs comprise large subfamilies (Hasin et al., 2014) that complicate evaluating their roles in biological processes. Thus, we turned to turnover-defective versions of Cdc42p to determine their impact on MAPK pathway activity. Most cells expressing Cdc42p^Q61L+TD^ resembled filaments (**Fig. 5B**, except for ∼ 16% of multi-budded cells). The filamentous morphology might result from Cdc42p- and fMAPK-dependent induction of genes that delay cell-cycle progression, resulting in elongated filamentous-type cells (Madhani et al., 1999). Thus, failure to turnover Cdc42p may stimulate the fMAPK pathway. In support of this possibility, the filamentous morphologies required an intact fMAPK pathway (**Fig. 5B**, *ste11Δ*). We also looked at the effect of Cdc42p turnover on mating. The formation of haloes of growth-arrested cells, which is induced by the mating pheromone *α*-factor, was not influenced by Cdc42p^Q61L+TD^ (**Fig. 5B**, bottom panel). This result indicates that turnover of Cdc42p impacts the fMAPK pathway but not the mating pathway.

**Figure 5.**
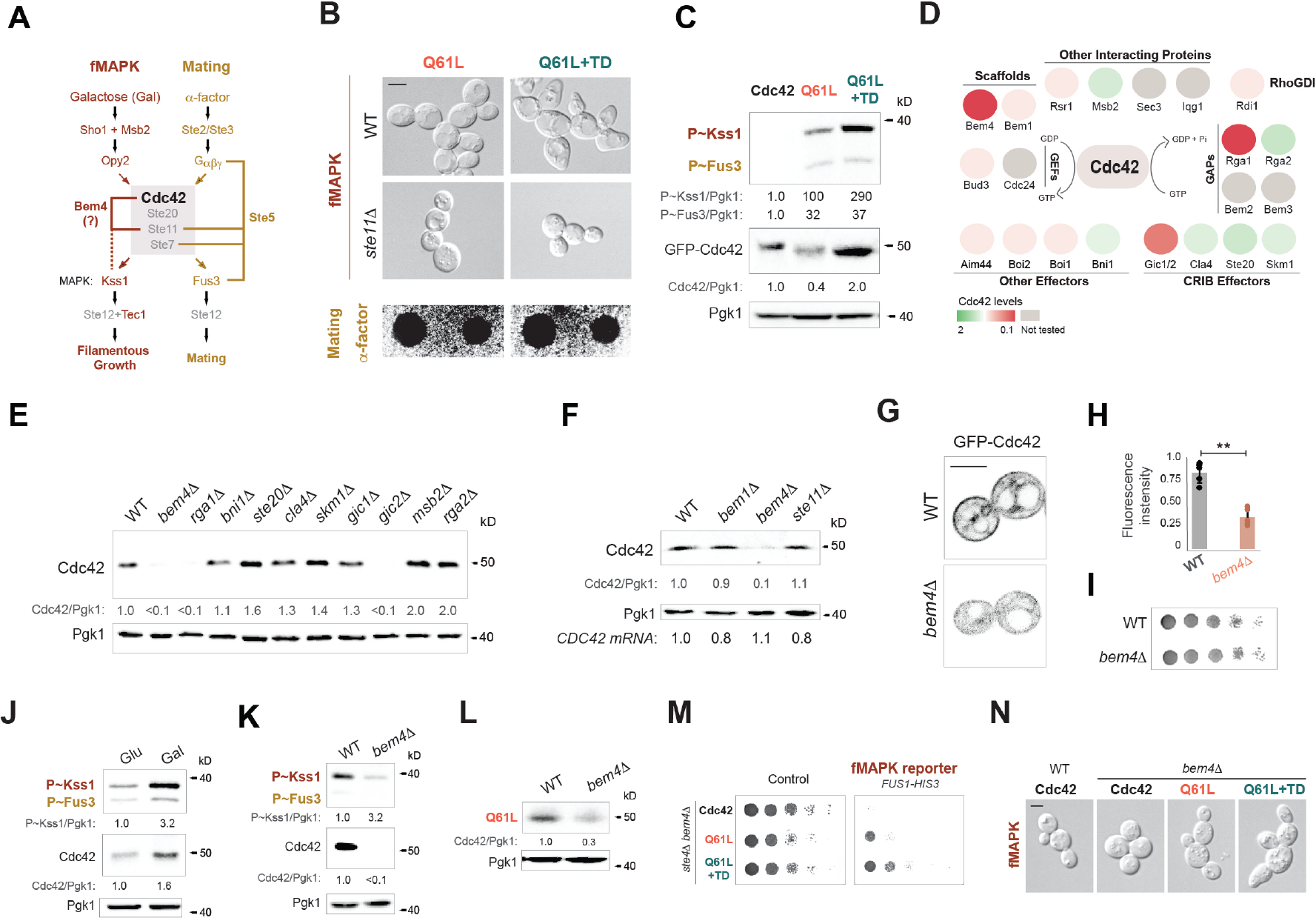
Turnover of Cdc42p differentially impacts MAPK pathway signaling: role of the scaffold Bem4p in stabilization of the Cdc42p protein. **A)** The model represents two Cdc42p-depenedent MAP kinase pathways. Colored text refers to fMAPK- (red) and mating-(yellow) specific components. Grey, common or shared factors. How Bem4p induces a pathway-specific signal at the level of Cdc42p remains unclear (question mark). **B)** Top, WT cells and *ste11Δ* expressing GFP-Cdc42p^Q61L^ (Q61L) or GFP-Cdc42p^Q61L,K5,94,96R^ (Q61L+TD). Bottom, halo formation in response to *α*-factor of WT cells expressing pGFP-Cdc42p^Q61L^ (Q61L) or pGFP-Cdc42^Q61L+TD^ (Q61L+TD). Cells were spotted onto SD-URA media, and *α*-factor was spotted at two concentrations, 6 µM or 2 µM. **C)** WT cells expressing GFP-Cdc42p (Cdc42), GFP-Cdc42p^Q61L^ (Q61L) or GFP-Cdc42p^Q61L,K5,94,96R^ (Q61L+TD) were grown for 6 h in SD-URA media. Numbers indicate relative P∼Kss1p, P∼Fus3p and GFP-Cdc42p compared to Pgk1p. **D)** Diagram of Cdc42p-interacting proteins. Colors indicate Cdc42p levels in extracts prepared from cells lacking the indicated protein. **E)** Cdc42p levels in WT cells and indicated mutants. Numbers indicate relative protein quantification. Cdc42p levels in cells lacking Gic2p and Bem4p were verified in multiple strains (*Fig. S5A*). **F)** Cdc42p levels in WT cells and *bem1Δ*, *bem4Δ*, *ste11Δ* mutants using antibodies to Cdc42p. Numbers indicate relative protein quantification. *CDC42 mRNA* levels were detected by RT-PCR. **G)** WT cells and *bem4Δ* mutant expressing GFP-Cdc42p were grown for 4 h at 30°C and examined by fluorescence microscopy. Scale bar, 5 µm. **H)** Relative fluorescence intensity of GFP-Cdc42p (Cdc42) or GFP-Cdc42p^Q61L^ (Q61L) in WT cells and the *bem4Δ* mutant. n >25, asterisk p-value <0.01. **I)** Serial dilutions of WT and *bem4Δ* cells expressing GFP-Cdc42p on SD-URA media. **J)** Analysis of P∼Kss1p, Cdc42p and Pgk1p levels in cells grown in SD (Glu) and in YEP-GAL (Gal) media for 6 h. **K)** WT and *bem4Δ* cells were grown for 4 h in YEPD and analyzed and described in panel 5J. **L)** GFP-Cdc42^Q61L^ levels in WT cells and *bem4Δ* mutant. **M)** Serial dilutions of the *ste4Δ bem4Δ* double mutant containing the growth reporter *FUS1-HIS3* and expressing the indicated alleles of Cdc42p were grown on SD-URA plates (control) and SD-URA-HIS + 1.6µM ATA (fMAPK reporter). **N)** Microscopic examination of WT cells and *bem4Δ* mutant expressing GFP-Cdc42p (Cdc42), GFP-Cdc42p^Q61L^ (Q61L) or GFP-Cdc42p^Q61L,K5,94,96R^ (Q61L+TD) grown for 6 h in SD-URA media.

The fMAPK and mating pathways are composed of distinct MAP kinases that regulate different transcription factor complexes to induce specific target genes and responses. The phosphorylation of the MAP kinase for the fMAPK pathway (Kss1p) and mating pathway (Fus3p) were examined, which provides a diagnostic readout of MAPK pathway activity (**Fig. 5C**). Compared to wild-type Cdc42p, expression of Cdc42p^Q61L^ stimulated the fMAPK and mating pathways, although it stimulated the fMAPK pathway to higher levels (**Fig. 5C**, ∼ 3-fold more than the mating pathway). This may be because Fus3p exists in a conformationally inactive state in the absence of pheromone and cannot be activated without binding to the scaffold Ste5p (Good et al., 2009). Cdc42p^Q61L+TD^ further stimulated the fMAPK pathway (**Fig. 5C**, by 2.9-fold more than Q61L alone) but did not impact the activity of the mating pathway. Therefore, activation of Cdc42p and accumulation of the active form of the protein result in preferential activation of the fMAPK pathway. As far as we are aware, this is the first case for Rho GTPase turnover influencing MAPK pathway signaling, which in this case results in a pathway-specific output.

During bud emergence and MAPK signaling, Cdc42p interacts with different proteins to execute an array of biological functions. Proteins that associate with Cdc42p may influence the stability and/or turnover of the protein. Cdc42p levels were examined by IB analysis in cells lacking Cdc42p-interacting proteins. Among a panel of mutants tested in the *Σ*1278b background, Cdc42p levels were found to be reduced in cells lacking Bem4p, Rga1p, and Gic2p (**Fig. 5**, **D** and **E**; *Fig. S5A*). Based on the results shown above (*Fig. S1A,* and *B*), lower levels of Cdc42p might be expected in the *rga1Δ* mutant, because cells lacking this GAP show elevated levels of GTP-bound Cdc42p (Gladfelter et al., 2002; Marquitz et al., 2002; Smith et al., 2002). However, the increase in GTP-Cdc42p in the *rga1Δ* mutant is modest (Smith et al., 2002), which may indicate that Rga1p impacts Cdc42p levels by multiple mechanisms. A role for Bem4p in regulating Cdc42p levels was unexpected, and we verified that cells lacking Bem4p showed reduced levels of Cdc42p in multiple strains and backgrounds (*Fig. S5B*). Bem4p has homology with members of the SmgGDS family of proteins (Hamel et al., 2011; Jennings et al., 2018; Shimizu et al., 2018; Shimizu et al., 2017; Vikis et al., 2002; Vithalani et al., 1998) and functions as a pathway-specific scaffold for the fMAPK pathway (Basu et al., 2020; Pitoniak et al., 2015). Unlike Rga1p, Bem4p does not impact Cdc42p activity (Hirano et al., 1996; Mack et al., 1996; Pitoniak et al., 2015) and may influence Cdc42p levels in a different manner. We verified by IB analysis (**Fig. 5**, E and F, 10-fold; *Fig. S5C*) and fluorescence microscopy (**Fig. 5, G** and **H** 4.2-fold) that Bem4p was required for cells to produce normal levels of Cdc42p. By comparison, loss of another Bem-type adaptor for Cdc42p, Bem1p (Basu et al., 2020; Bender and Pringle, 1991; Butty et al., 2002; Irazoqui et al., 2003; Park et al., 1997; Pitoniak et al., 2015), did not impact Cdc42p levels (**Fig. 5F**). Bem4p did not impact *CDC42* gene expression (**Fig. 5F**, *CDC42* mRNA), and Cdc42p levels were not impacted in cells lacking an intact fMAPK pathway (**Fig. 5F**, *ste11Δ*), which indicates that Bem4p regulates Cdc42p levels separate from its role in regulating the fMAPK pathway. Bem4p did not impact the levels of a membrane-anchored version of GFP, GFP^KKSKKCATL^ (*Fig. S5D*). Given that Bem4p interacts with Cdc42p *in vitro* and *in vivo* (Drees et al., 2001; Hirano et al., 1996; Mack et al., 1996; Pitoniak et al., 2015), Bem4p may stabilize Cdc42p levels by binding to the Cdc42p protein.

Cells lacking Bem4p did not exhibit polarity (**Fig. 5G**) or growth defects (**Fig. 5I**). Thus, cells produce more Cdc42p than is necessary for growth. This idea is already appreciated: for example, cells expressing the *cdc42-1* allele show low levels of Cdc42p yet are viable at permissive temperatures (Adamo et al., 2001). Excess Cdc42p levels might have pathway-specific consequences. Specifically, by stabilizing the Cdc42p protein, Bem4p may selectively promote activation of the fMAPK pathway. Cells grown in the non-preferred carbon source, galactose, which induces filamentous growth and fMAPK pathway activity, showed higher levels of Cdc42p (**Fig. 5J**, Cdc42), which corresponded to elevated phosphorylation of Kss1p (**Fig. 5J**, P∼Kss1p). The induction of the fMAPK pathway and the elevated levels of Cdc42p that occurred during growth in galactose were dependent on Bem4p (**Fig. 5K**). Bem4p also stabilized the levels of the GTP-locked version, Cdc42p^Q61L^ (**Fig. 5L**). Stabilization of GTP-bound Cdc42p would be expected to result in elevated fMAPK pathway activity by activation of the effector PAK Ste20p.

Bem4p binds to other proteins to regulate the fMAPK pathway, including the GEF Cdc24p and the MAPKKK, Ste11p (Pitoniak et al., 2015). To determine the contribution that Bem4p-dependent stabilization of Cdc42p plays on the activity of the fMAPK pathway, Cdc42p^Q61L+TD^ was expressed in cells lacking Bem4p. In the *bem4Δ* mutant, Cdc42p^Q61L+TD^ restored fMAPK pathway activity based on a transcriptional growth reporter (**Fig. 5M**). By comparison, Cdc42p^Q61L+TD^ was unable to bypass the signaling defect of cells lacking the MAPKKK Ste11p (*Fig. S5E*), which propagates signals initiated by Ste20p. Cdc42p^Q61L+TD^ also restored filamentous growth to the *bem4Δ* mutant (**Fig. 5N**). Therefore, stabilization of Cdc42p by Bem4p is critical for Cdc42p function in the fMAPK pathway.

## Discussion

We show here that the yeast Rho GTPase Cdc42p, one of the most highly studied and best-understood members of the Rho GTPase family, is regulated by turnover. This is an important discovery that will permit studying Rho GTPase turnover regulation in a genetically-tractable model system. Using this system, we show that Cdc42p is ubiquitinated and turned over in the proteasome by the NEDD4-type E3 ubiquitin ligase Rsp5p. We also show that versions of the protein that mimic the active conformation of Cdc42p are preferentially turned over compared to the wild-type species, which provides a mechanism for modulating the activity of this evolutionarily conserved regulator of cell polarity and signaling.

By exploring the mechanism of Rsp5p-dependent turnover of Cdc42p, we discovered a role for Hsp40p and Hsp70p chaperones in regulating Cdc42p degradation. This discovery extends previous findings linking HSPs to Rsp5p-dependent turnover of misfolded proteins (Fang et al., 2014). HSPs are global regulators of cell polarity and signal-dependent cell differentiation (Calderwood and Gong, 2016), and evolutionary drivers of morphogenetic responses and phenotypic plasticity (Rutherford and Lindquist, 1998). Our findings provide the first connection between HSPs and Rho GTPase turnover. HSPs may regulate Rho GTPase folding and turnover to control Rho-driven biological processes. For example, Hsp40p and Hsp70p chaperones regulate cell-cycle progression and cell size (Ferrezuelo et al., 2012; Truman et al., 2012; Vergés et al., 2007), which are connected to Cdc42p-dependent morphogenesis. Remarkably little is known about the folding of Rho GTPases, and HSPs may provide a mechanism for Rho GTPase protein quality control. Cdc42p is turned over rapidly at high temperatures and can form aggregates when overexpressed or when versions of the protein are expressed that compromise the stability of the protein (**Fig. 6A**). To our knowledge, this is the first case of aggregate formation by a Rho GTPase. HSPs regulate turnover of cytosolic proteins to prevent protein aggregation (Amm et al., 2014). Given that Cdc42p aggregates preferentially accumulate in mother cells, HSP-dependent turnover of Cdc42p and other Rho GTPases may impact aging, polarity health, and the rejuvenation of cell polarity. In humans, elevated CDC42 levels impact aging and can induce higher mortality (Kerber et al., 2009). CDC42 levels also regulate senescence (Umbayev et al., 2018) and aging in in stem cells (Florian et al., 2012). Rho GTPase turnover by HSPs may be a general mechanism that has far-reaching implications in human health and disease.

**Figure 6.**
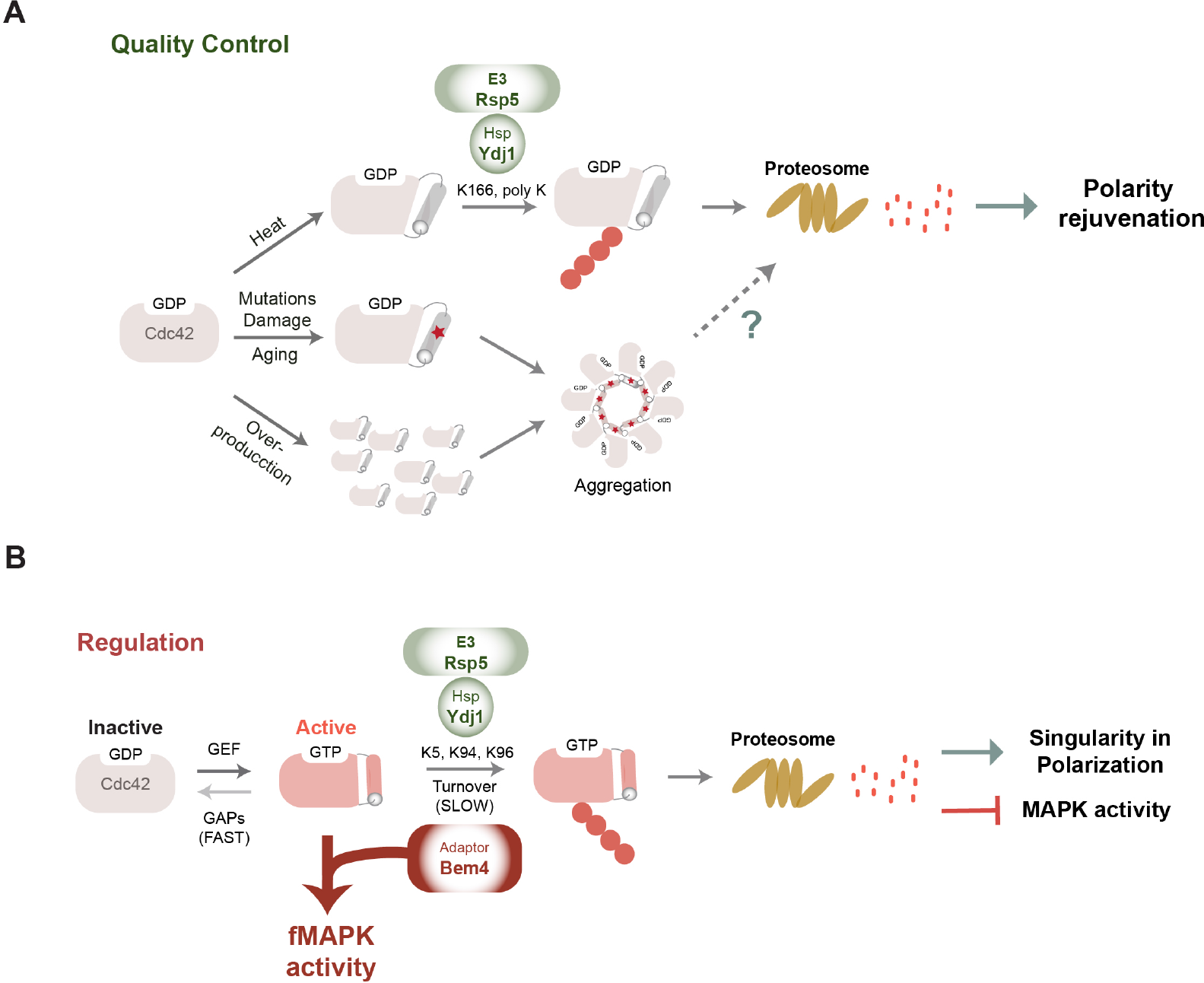
Cdc42p turnover regulation by HSPs and Rsp5p impacts quality control, as well as singularity in polarization, MAPK signaling and polarity rejuvenation. **A)** Cdc42p turnover occurs by HSPs as part of a quality-control mechanism. Elevated temperatures (Heat) stimulate Cdc42p turnover, which occurs through Ydj1p, Rsp5p, and the proteasome. Overexpression of the protein, or elevated levels resulting from failure of the protein to be turned over, such as in cells lacking Ydj1p, lead to aggregation of the protein that can stimulate turnover. Aggregation can result from mutations or protein damage (star) that leads to instability. Such problems can accrue during cellular aging; hence, we speculate that the turnover of mis-folded Cdc42p can promote the overall ‘polarity health’ of cells and lead to the rejuvenation of cell polarity. Question mark, aspects of the turnover of Cdc42p aggregates are not clear. **B)** Cdc42p is activated by GEFs and inactivated by GAPs, which is thought to occur rapidly (FAST). Here we show that yeast Cdc42p is turned over by a mechanism that involves the HSP chaperone Ydj1p and the E3 ubiquitin ligase Rsp5p, which is slower than GAP hydrolysis (SLOW). Ubiquitin conjugates of Cdc42p (large red dots) results in degradation of the protein in the proteasome. Conformational changes that occur to the protein when activated by binding to GTP may lead to accelerated turnover of the protein by recognition by Ydj1p and/or Rsp5p. Turnover of Cdc42p promotes singularity in polarization, and is expected to attenuate MAPK pathway signaling (block arrow). Bem4p is a Cdc42p-interacting protein that may function to stabilize Cdc42p (red arrow). As a result, one MAPK pathway is stimulated among other Cdc42p-dependent MAPK pathways in the cell.

Remarkably, Hsp40p promoted turnover of the GTP-locked versions of Cdc42p, thereby implicating HSP function to the regulation of Rho GTPase activity (**Fig. 6B**). Different lysines were required for the turnover of Cdc42p in response to temperature and its activation state (**Fig. 6**), and it will be interesting to explore similarities and differences between these related regulatory mechanisms. The GTP-bound conformation of Cdc42p may be recognized by HSPs as the preferred species due to changes in the folding of the protein. Alternatively, HSPs may degrade GTP-bound Cdc42p molecules that mis-fold over time. In addition, the GTP-bound conformation is preferentially localized to the plasma membrane, a location where Rsp5p recognizes substrate proteins.

The mechanism by which Cdc42p induces a growth site is well defined (Caviston et al., 2002; Irazoqui et al., 2003; Slaughter et al., 2009; Woods et al., 2016). How cell polarity is attenuated has also been extensively studied, and it is clear that negative feedback imparts robustness to polarity establishment (Howell et al., 2012; Kuo et al., 2014; Ozbudak et al., 2005). An interesting feature of polarity regulation is the competition among growth sites (Howell et al., 2012; Wu et al., 2015). How competition is resolved among competing sites is less clear. We show that versions of active Cdc42p that cannot be turned over make multiple buds. This might indicate that turnover of Cdc42p helps to resolve competition at sites where Cdc42p polarized to ensure singularity in cell polarization. Alternatively, the turnover of active Cdc42p might impact polarity establishment through another mechanism that can be elucidated upon further investigation.

GAPs promote the rapid (within a few seconds) hydrolysis of GTP to GDP (Freisinger et al., 2013). By comparison, the turnover of Cdc42p may be slow (half life < 30min for Q61L). Therefore, turnover might represent a slow and more permanent way to remove active Cdc42p (**Fig. 6B**, see fast and slow). Moreover, GAPS and turnover play different roles in Cdc42p regulation. As cells lacking multiple GAPs (*rga1Δ rga2Δ bem3Δ*) exhibit polarity defects but do not produce cell with multiple buds (Caviston et al., 2003; Smith et al., 2002). It will be interesting to explore the different roles GAPs and turnover play in modulating Cdc42p function. Generally speaking, the discovery of Cdc42p turnover may deepen our understanding of cell polarity regulation in yeast and other systems.

The turnover regulation of Cdc42p also induces pathway-specific effects on Rho GTPase function. In particular, stabilization of Cdc42p by the fMAPK pathway-specific adaptor, Bem4p, leads to differential activation of the fMAPK pathway (**Fig. 6B**). This discovery is important because it provides the first link between Rho GTPase turnover/stability and MAPK pathway regulation. Bem4p, itself related to members of the SmgGDS family of proteins, may function as a chaperone to promote folding of Cdc42p. Alternatively, Bem4p may prevent accessibility of HSPs and Rsp5p by binding to Cdc42p and blocking sites of ubiquitination. This mechanism is contingent upon the fact that the MAP kinase for the mating pathway, Fus3p, is conformationally closed until the mating pathway is activated (Good et al., 2009). Therefore, stabilizing GTP-bound Cdc42p may preferentially induce the fMAPK Kss1p, which is not conformationally constrained. Once activated, the fMAPK pathway is attenuated, in part, by ubiquitin-dependent turnover of the mucin-type sensor, Msb2p (Adhikari et al. 2015). Msb2p is also proteolytically processed to activate the pathway by release of an auto-inhibitory domain (Vadaie et al., 2008). Thus, a balance of proteolytic functions shapes the kinetics of the filamentation response. Amplification of signaling by Rho GTPase stabilization may be a general mechanism for directing these proteins to pathway-specific contexts. Intriguingly, in fungal pathogens, Cdc42p and other Rho GTPases contribute to many fungal attributes underlying virulence (Brand et al., 2014; Chen et al., 2019; Silva et al., 2019). Intriguingly, HSPs have also been implicated in the regulation of fungal pathogenesis (Horianopoulos and Kronstad, 2021). It will be interesting to explore whether HSPs regulate Rho GTPase levels to control aspect of the pathogenic response.

## Materials and Methods

### Yeast strains, reagents and media

Strains are listed in *Table S1*. Plasmids are listed in *Table S2*. Primers are listed in *Table S3*. Yeast cultivation was performed in synthetic media (SD; 0.67% yeast nitrogen base without amino acids, 2% dextrose), supplemented with amino acids as required, <u>Y</u>east <u>E</u>xtract <u>P</u>eptone <u>D</u>extrose media [YEPD (1% bacto-yeast extract, 2% bacto-peptone, 2% dextrose)] and YEPGAL (1% bacto-yeast extract, 2% bacto-peptone, 2% galactose) at 30°C or indicated temperatures. YlpLac211-P*CDC42*-GFP-linker-*CDC42* plasmid was provided by the Lew Lab (Woods et al., 2016). The linker (GSCGRAPPRRLVHP) between GFP and Cdc42p improves functionality of the protein. To construct pRS316-GFP-linker-*CDC42P* (PC6454; pGFP-Cdc42), GFP-linker-*CDC42* (with *CDC42* promoter) was subcloned with *EcoR*I and *Sal*I into pRS316 [CEN/URA (Sikorski and Hieter, 1989)]. To create pRS316-PGAL1-GFP-linker-*CDC42* (p-GAL1-GFP-Cdc42), the cassette pGAL containing *KanMX6* antibiotic resistance marker from (Longtine et al., 1998) was generated by polymerase chain reaction (PCR) and inserted in the pGFP-Cdc42 plasmid by homologous recombination.

To insert point mutations in plasmids carrying *CDC42*, GeneART^TM^ Site-Directed Mutagenesis (SDM) Kit (Cat#A13282, Thermo Fisher) was used according to the manufacturer’s protocols. Nucleotides corresponding to lysines were mutated in pairs or groups because non-preferred lysines can be ubiquitinated when a preferred lysine is absent (Kravtsova-Ivantsiv, 2011). Some mutations were introduced by *in vivo* homologous recombination in yeast. Briefly, primers containing the desired point mutations and a flanking region for homologous recombination were amplified by PCR using pGFP-Cdc42 as template. pGFP-Cdc42 was linearized by PshAI (Cat#R0593S, New England Biolabs) and co-transformed with the PCR product into an uracil auxotrophic strain. Plasmid rescue was performed from colonies arising on SD-URA semi-solid agar media. Nucleotide sequences were confirmed by Sanger Sequencing (Genewiz). A similar approach was used to generate p-GFP-linker-KKSKKCTIL. In this case the plasmid pGFP-Cdc42 was linearized with the restriction enzyme PshAI and primers containing the flanking region for homologous recombination between the linker and the 3’-end of the sequence of *CDC42* were used. To generate the pHis6x-linker-Cdc42 and pHis6x-linker-Cdc42^13KR^ by homologous recombination, pGFP-Cdc42 was linearized with the restriction enzyme SnaBI (Cat#R0130S, New England Biolabs), and co-transformed in yeast with primers harboring the *HISX6* sequence and flanking regions to the *CDC42* promoter and linker, sequences listed in *Table S3*. Gene disruptions were performed by antibiotic resistance markers *NAT*, *HYG* and *KanMX6* using PCR-based approaches using published templates (Goldstein and McCusker, 1999; Longtine et al., 1998).

### Quantitative Polymerase Chain Reaction (qPCR) Analysis

qPCR analysis was performed as described (Prabhakar et al., 2020). 0.02 O.D. of yeast cells, WT (PC538), *bem1Δ* (PC6680), *bem4Δ* (PC3351), *ste11Δ* (PC6604) were inoculated in SD media. Samples were grown at 30°C and collected after 4 h. RNA extraction was performed by hot acid phenol-chloroform followed by a purification step with the RNeasy Mini Kit (Cat#79254). RNA stability was determined by agarose gel electrophoresis in 1.2 % agarose Tris-Borate-EDTA (TBE, 89 mM Tris base, 89 mM Boric acid, 2mM EDTA). For reverse transcription reactions, RNA concentration was adjusted to 60 ng/μL. Reverse transcription was performed iScript Reverse Transcriptase Supermix (BioRad, 1708840). qPCR was performed using iTaq Universal SYBR Green Supermix (BioRad, 1725120) following the manufacturer’s instructions. Reactions contained 10 μl samples (180 ng/μL cDNA, 0.2 μM each primer, 5 μl SYBRGreen master mix). qPCR was performed using an BioRad thermocycler (CFX384 Real-Time System; Applied Biosystems). Relative gene expression was calculated using the 2^−Δ**Ct**^ formula, where Ct is defined as the cycle at which fluorescence was determined to be statistically significant above background; ΔCt is the difference in Ct of the *CDC42* gene and housekeeping gene (*ACT1*). The primers used are listed in *Table S3*. Values represent the mean of at least two independent biological replicates and two technical replicates.

### Protein turnover

CHX assays were performed as described in (Adhikari et al., 2015a). 100 ml of wild-type cells (PC538) and cells expressing pGFP-Cdc42 (PC6454) or pGFP-Cdc42^Q61L^ (PC7458) at 0.02 of O.D. were grown in SD or SD-URA (to maintain plasmid selection) for 4 h. After 4 h, the medium was supplemented with 25 μg/ml of CHX, and 10 mL of samples were collected at 0, 15, 30, 45, 60, 90 and 120 min to generate cell extracts for IB analysis. Experiments were performed in two independent biological replicates. For pGAL1-promoter shutoff experiments were done based on (Adhikari et al., 2015b). Wild-type cells expressing the pGAL1-GFP-Cdc42 (PC7349) were grown in YEPGAL (1% bacto-yeast extract, 2% bacto-peptone, 2% galactose) media for 4 h and transferred to YEPD media. 10 ml of samples were collected at 0, 60, 90 and 120 min. Experiments were performed in three independent biological replicates.

The proteasome inhibitor MG132 (carbobenzoxy-Leu-Leu-leucinal; CAS 133407-82-6, Calbiochem) was reconstituted with ethanol (20 mg/ml) according to the manufacturer’s protocol. 20 mL of wild-type cells containing GFP-Cdc42p or GFP-Cdc42p^Q61L^ at 0.02 O.D. were grown at 30°C for 4 h in SD-URA. At 4 h, media was supplemented with 0.5% ethanol (control) or 75μM MG132 and cells were harvested after 2 h to generate extracts for IB analysis. Experiments were done in two independent replicates.

### Co-IPT Analysis

Cells for Co-IPT analysis were grown and lysed as previously reported (Adhikari et al., 2015a). Wild-type cells grown to mid log phase in SD media or cells expressing the pGFP-Cdc42 or pGFP-Cdc42^Q61L^ grown to mid log phase in SD-URA media. Cells were harvested by centrifugation, washed with 1% phosphate -based saline (PBS) and resuspended in 500 µl of IP buffer (50mM Tris-Cl, pH 8, 1mM EDTA, 100mM NaCl, 1.5% NP-40, 1 mM phenylmethanesulfonyl fluoride (PMSF), 1x protease inhibitor cocktail). 200 µl of glass beads were added, and the cells were lysed by vortexing (FastPrep-24, MP Biomedicals) for 10 min at 4°C. After a 30 min incubation at 4°C, lysed cells were centrifuged at 4°C for 10 min at 14,000 rpm. The immunoprecipitation was performed as described in (Elu et al., 2019). Briefly, clarified lysate was incubated with 25 µl GFP-Trap magnetic beads (ChromoTek GFP-Trap^®^ Magnetic Agarose), at 4°C for 2 h. Beads were separated by a magnet (Cat#1614916, Bio-Rad, Inc.) and washed twice with 500 µl of dilution buffer (10 mM Tris-Cl pH 7.5; 150 mM NaCl; 0.5 mM EDTA). Proteins were eluted from the beads by treatment at 95°C for 2 h in 2 x Sodium Dodecyl Sulfate-polyacrylamide (SDS)-sample buffer (120 mM Tris-Cl pH 6.8; 20 % glycerol; 4% SDS; 0.04 % bromophenol blue; 10 % BME). The supernatant was examined by IB analysis.

### Immunoblot Analysis

Cells were grown to saturation in SD or YEPD media for 16 h and transferred fresh media and grown for 4-6 h to mid log phase. Cells were harvested by centrifugation. Proteins extracts were prepared by mechanical disruption with beads followed by a trichloroacetic acid (TCA) precipitation method (Basu et al., 2020). Protein precipitates were analyzed by SDS-PAGE and transferred to a nitrocellulose membrane (Cat#10600003, Amersham^TM^ Protran^TM^ Premium 0.45 μm NC, GE Healthcare Life sciences). Monoclonal mouse anti-GFP antibodies were used (Cat#11814460001, clones 7.1 and 13.1, Roche) at 1:1,000 dilution. Polyclonal rabbit phospho-p44/42 MAPK (Erk1/2, Cat#3102, Cell Signaling Technology) were used at 1:10,000 dilution. Mouse anti-Kss1p antibodies (yC-19, Santa Cruz Biotechnology) were used at 1:10,000 dilution. Monoclonal mouse anti-ubiquitin antibodies were used at 1:5,000 dilution (P4G7, Santa Cruz Biotechnology). Rabbit anti-Cdc42 antibodies (Kozminski et al., 2000) were used at 1:1,000 dilution and were generously provided by Dr. Keith Kozminski (University of Virginia). Monoclonal mouse anti-Pgk1 antibodies (22C5D8, Cat#459250, Invitrogen) were 1:1,000 dilution. Secondary anti-mouse IgG-HRP (Cat# 1706516, Bio-Rad Laboratories) and goat anti-rabbit IgG-HRP (Cat#115-035-003, Jackson ImmnunoResearch Laboratories) were used. The nitrocellulose membrane was blocked with 5% non-fat dried milk or 5% bovine serum albumin (BSA) (BSA; for p44/42 antibody) for 1 h prior antibody detection. Primary incubations were performed at 4°C for 16 h and secondary at 20°C for 1 h. IBs were visualized by Gel Doc XR Imaging System (Bio-Rad, Inc.), after addition of Chemiluminescent HRP substrate for chemiluminescent Westerns (Radiance^TM^ Plus Substrate, Azure Biosystems).

Band intensities quantitation of P∼Fus3p, P∼Kss1p, Cdc42p, GFP-Cdc42p and ubiquitin were detected under non-saturated conditions and normalized to the housekeeping protein Pgk1p using the Image Lab Software (Bio-Rad, Inc.). Wild-type cells and control conditions were set to 1 and adjusted for other samples accordingly.

### fMAPK Reporter

The activity of the fMAPK pathway was tested by using *FUS1-HIS3* growth reporter (McCaffrey et al., 1987). Cells lacking an intact mating pathway (*ste4*) mating, show basal activity of the fMAPK pathway by this reporter (Cullen et al., 2004). Wild-type cells (PC538) and a control strain (*ste11Δ*, PC3862), were growth in SD-URA media to maintain plasmid selection (SD-URA-HIS, control), and media lacking Histidine (SD-URA-HIS) or supplemented with ATA (3-amino-1,2,4-triazole) to evaluate fMAPK activity.

### Fluorescence microscopy

The localization of Cdc42p was examined in strains containing the plasmid pGFP-Cdc42, in this plasmid the expression of the GFP-tagged Cdc42p is controlled by its own promoter. In all experiments, cells were grown to mid log phase at 30°C in SD-URA media before examination, except in the case of the temperature sensitive mutants, *rsp5-1* and *cim3-1*, which were additionally grown at 37°C for 2 h. Differential interference contrast (DIC), fluorescence microscopy using fluorescein isothiocyanate (FITC) and Rhodamine filter sets were used in an Axioplan 2 fluorescence microscope (Zeiss) with a Paln-Apochromat 100x/1.4 (oil) objective with the Axiocam MRm camera (Zeiss). Images were analyzed using Axiovision 4.4 software (Zeiss). Actin stanning was performed as described (Amberg et al., 2006a) using Phalloidin-Atto 532 (Millpore Sigma, MA, 49429). Vacuoles and endocytic compartment were stained with the lipophilic styryl dye FM4-64 [(*N*-(3-Triethylammonumporpyl)-4-(6-(4-(Diethylamini) Phenyl) Hexatrienyl) Pyridinium Dibromide); Cat#T13320, ThermoFisher) as described in (Amberg et al., 2006b), for GFP-Cdc42p overexpression cells carrying the pGAL1-GFP-Cdc42 were grown in YEPGAL media for 4 h.

Images were analyzed in Adobe Photoshop and ImageJ. Fluorescence images were converted to Grayscale and inverted using ImageJ. For fluorescence intensity quantification, the corrected total fluorescence (CTCF) was determined by using the measure tool of the software ImageJ. To calculate CTCF, the area selected multiplied by fluorescence background was subtracted to the integrated density (Keith R. Porter Imaging Facility, UMBC, Baltimore). Images taken with the same exposure time were compared, and in all analysis >4 frames containing >10 cells were used. For relative quantification the highest value of the control condition was set to 1 and the other values were calculated accordingly. Area of Cdc42p aggregates was calculated by using the measure tool of ImageJ from two independent replicates.

### Time-Lapse Microscopy

Experiments were performed based on (Prabhakar et al., 2020). Cells were grown at 30°C for 16 h in SD-URA. 5 μL of cells diluted to 0.02 O.D. were placed under 1% agarose pads using a 12 mm Nunc glass base dish (Cat#150680, Thermo Scientific, Waltham). A wet cotton pad was placed around the agar to prevent dehydration. Cells were grown at 30°C for 2 h prior to imaging. Live-cell microscopy was performed with a Zeiss 170 confocal microscope equipped with a Plan-Apochromat 40x/1.4 Oil DIC M27 objective. During imaging Cdc3-mCherry cells (PC7365) expressing different alleles of GFP-Cdc42p were grown at 30°C for 4 h and images were taken in intervals of 10 min. For the detection of GFP-Cdc42p a 488nm laser (496nm-548nm filter), and for Cdc3-mCherry, a 580 nm laser (589nm-708nm filter) were used. Images were taken with multiple Z-stacks (8-10) at 1 μm increments. Images were analyzed with ImageJ (https://imagej.nih.gov) using the Z-project and template matching plugins.

### Statistical analysis

All statistical analysis were performed with the 2021.1 XLSTAT software (https://www.xlstat.com). Student’s t-test was used to determine statistical significance and generate P-values.

#### Abbreviations

ATA: (3-amino-1,2,4-triazole)
BSA: bovine serum albumin
CFW: calcofluor white
CHX: cycloheximide
DIC: differential interference contrast
Glu: glucose
Gal: galactose
GAP: GTPase activating protein
GEF: guanine nucleotide exchange factor
GFP: green fluorescent protein
GTPase: guanine nucleotide triphosphate
IB: Immunoblots
IPT: Immunoprecipitation
HR: homologous recombination
HSP: heat shock protein
MAPK: mitogen-activated protein kinase
fMAPK: filamentous growth MAP kinase pathway
O.D.: optical density
PAK: p21-activated kinase
Rho: Ras homology
S.D.: standard deviation
SDM: site-directed mutagenesis
SDS-PAGE: sodium dodecyl sulfate-polyacrylamide gel electrophoresis
TD: turnover deficient
YEPD: yeast extract peptone and dextrose
WT: wild type

## Supporting information

suppemental Data

## Acknowledgments

Thanks to John Pringle (Stanford University), Charles Boone (University of Toronto), Keith Kozminski (University of Virginia, Charlottesville, VA), Daniel Lew (Duke University, Durham NC), Thibault Mayor (University of British Columbia, Vancouver, Canada), Chris Burd (Yale University, New Haven, CT), David Pellman (Harvard Medical School, Boston, MA), and Scott Emr (Cornell University, Ithaca, NY) for providing reagents. Thanks to Rick Cerione for reading the manuscript. Thanks to Aditi Prabhakar and Alan Seigel for assistance with confocal microscopy and lab members for suggestions. The work was supported by a grant from the NIH (GM098629).

