## Supplementary material for "Turnover Regulation of the Rho GTPase Cdc42 by Heat Shock Protein Chaperones and the MAPK Pathway Scaffold Bem4": suppemental Data

**MOVIES AND SUPPLEMENTAL FIGURES**

***Movie 1***

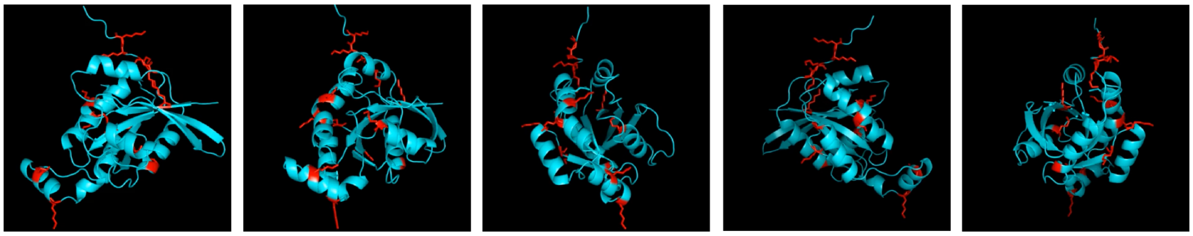

**Movie 1. Lysine residues of Cdc42p**. The yeast Cdc42p protein sequence was overlaid onto the crystal structure of human Cdc42p using the Expasy web server SWISS-MODEL (https://swissmodel.expasy.org) (Nassar et al. 1998). Side chains of lysine residues of Cdc42p yeast structure are colored in red.

***Movie 2***

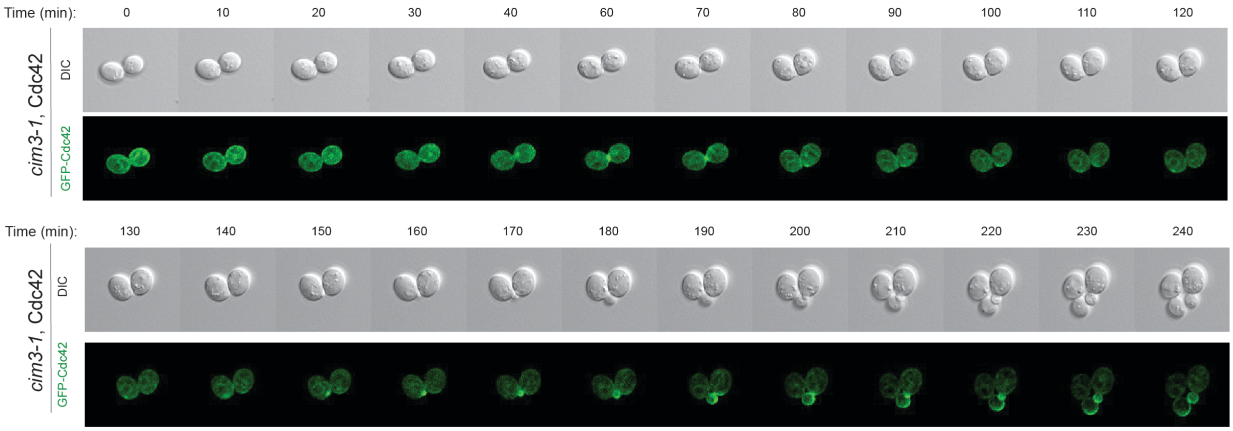


**Movie 2.** Confocal time-lapse microscopy of *cim3-1* (PC5852) cells expressing GFP-Cdc42p on SD-URA media grown at 36°C. Cells were grown for 3 h at 30°C. Cdc42p aggregates were observed in 2% of the cells. Time interval, 10 min.

#### Movie 3

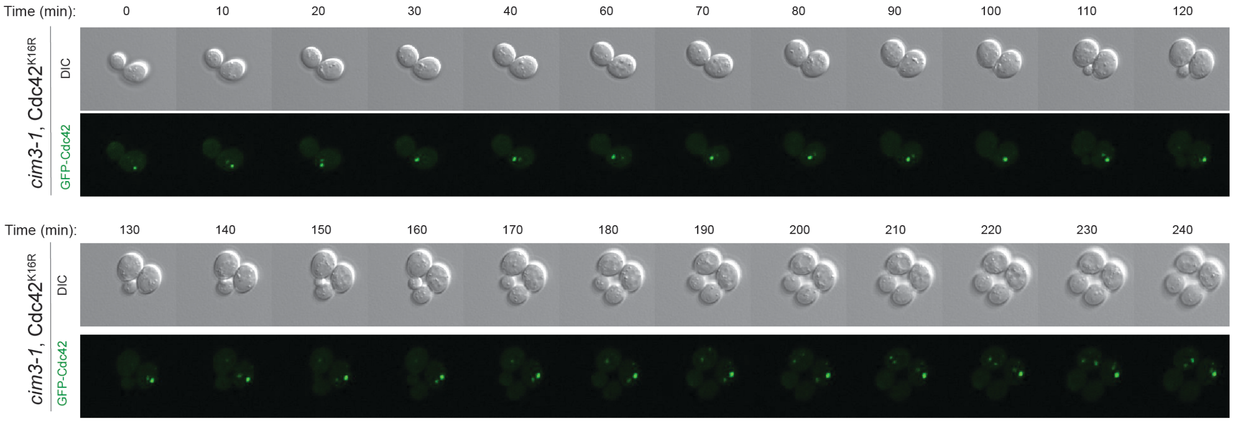


**Movie 3.** Confocal time-lapse microscopy of *cim3-1* (PC5852) cells expressing GFP-Cdc42p$^{K16R}$ on SD-URA media grown at 36°C. Cells were grown for 3 h at 30°C. Cdc42p aggregates were observed in 82% of the cells. Time interval, 10 min.

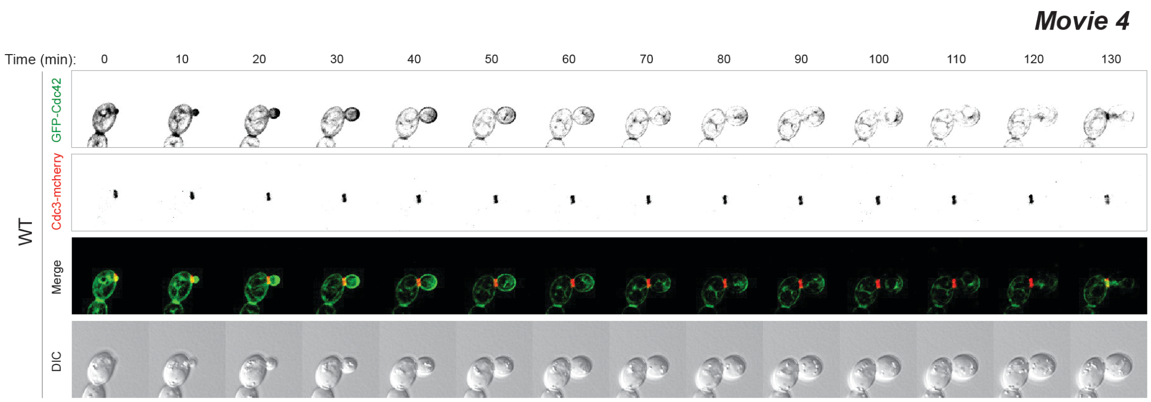


**Movie 4**. Confocal time-lapse microscopy of WT (PC7365) cells expressing GFP-Cdc42p and Cdc3-mcherry (septin marker) on SD-URA media. Cells were grown for 5 h at 30°C. Green refers to Cdc42p, and Cdc3-mcherry (red) is a septin ring marker. Time interval, 10 min.

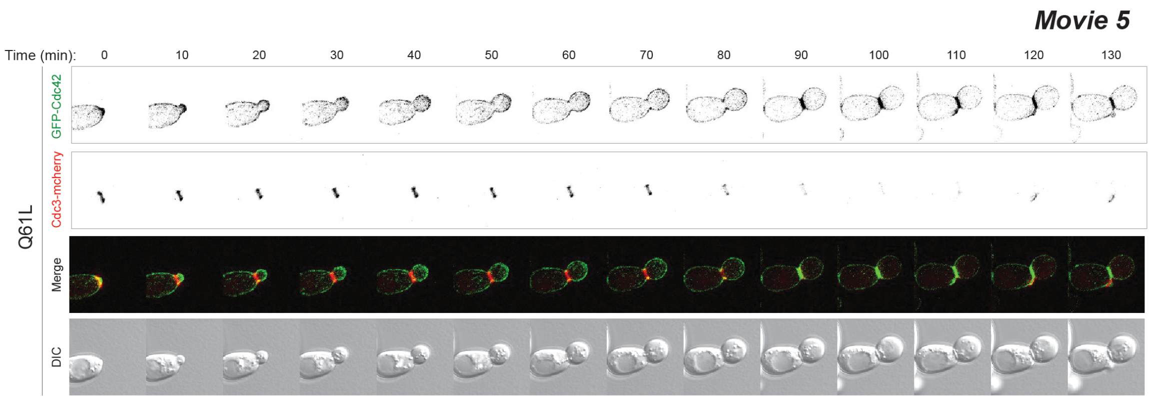


**Movie 5.** Confocal time-lapse microscopy of WT (PC7365) cells expressing GFP-Cdc42p$^{Q61L}$ and Cdc3-mcherry (septin marker) in SD-URA media. Cells were grown for 5 h at 30°C. Green refers to Cdc42p, and Cdc3-mcherry (red) is a septin ring marker. Time interval, 10 min.

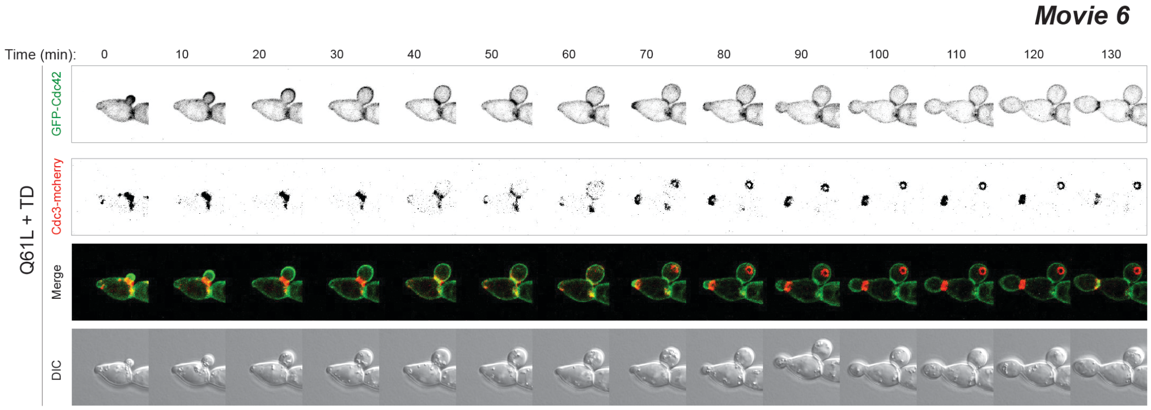


**Movie 6.** Confocal time-lapse microscopy of WT (PC7365) cells expressing GFP-Cdc42p$^{Q61L+K5,94,96R}$. and Cdc3-mcherry (septin marker) in SD-URA media. Cells were grown for 5 h at 30°C. Green refers to Cdc42p, and Cdc3-mcherry (red) is a septin ring marker. Time interval, 10 min.

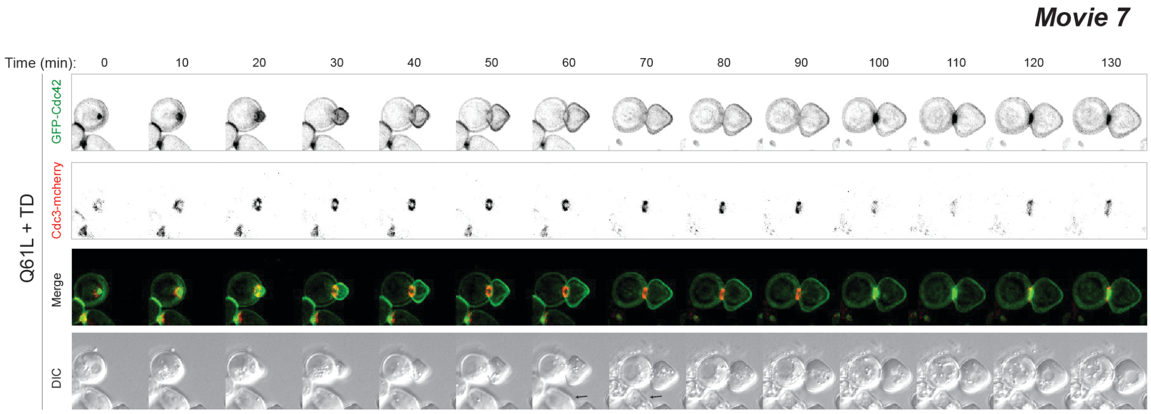


**Movie 7.** Confocal time-lapse microscopy of WT (PC7365) cells expressing GFP-Cdc42p$^{Q61L+K5,94,96R}$ and Cdc3-mcherry (septin marker) in SD-URA media. Cells were grown for 5 h at 30°C. Green refers to Cdc42p, and Cdc3-mcherry (red) is a septin ring marker. Time interval, 10 min.

### TABLES

#### Table S1. Yeast strains used in the study.

| Name | Genotype | Reference |
|---|---|---|
| PC313 | *MAT***a** *ura3-52* | (Liu et al. 1993) |
| PC538 | *MAT***a** *ste4 FUS1-lacZ FUS1-HIS3 ura3-52* | (Cullen et al. 2004) |
| PC673 | *MAT***a** *ste4 FUS1-lacZ FUS1-HIS3 ura3-52 ste20:: kanMX6* | (Cullen et al. 2004) |
| PC948 | *MAT***a** *ste4 FUS1-lacZ FUS1-HIS3 ura3-52 msb2:: kanMX6* | (Cullen et al. 2004) |
| PC986 | *S288c MATα his3Δ0, leu2Δ0, met15Δ0, ura3Δ0* | Research Genetics |
| PC969 | *MAT***a** *ste4 FUS1-lacZ FUS1-HIS3 ura3-52 cla4:: kanMX6* | (Pitoniak et al., 2015) |
| PC3063 | *S288c MATα his3Δ0, leu2Δ0, met15Δ0, ura3Δ0 pep4::kanMX6* | Research Genetics |
| PC3288 | *S288c* Y7092 *MATα ura3-52 his3-200 trp1-901 lys2-801 suc2-p leu2-3$^{e}$* | (Strochlic et al. 2008) |
| PC3290 | *S288c* Y7092 *MATα ura3-52 his3-200 trp1-901 lys2-801 suc2-p leu2-3$^{e}$ rsp5-1* | (Strochlic et al. 2008) |
| PC3391 | *MAT***a** *ste4 FUS1-lacZ FUS1-HIS3 ura3-52 rga1::NAT* | (Pitoniak et al. 2015) |
| PC3392 | *MAT***a** *ste4 FUS1-lacZ FUS1-HIS3 ura3-52 ste12::kanMX6 rga1::NAT* | (Pitoniak et al. 2015) |
| PC3393 | *MAT***a** *ste4 FUS1-lacZ FUS1-HIS3 ura3-52 bem4::HYG rga1::NAT* | (Pitoniak et al. 2015) |
| PC3551 | *MAT***a** *ste4 FUS1-lacZ FUS1-HIS3 ura3-52 leu2::HYG bem4::NAT* | (Pitoniak et al. 2015) |
| PC3862 | *MAT***a** *ste4 FUS1-lacZ FUS1-HIS3 ura3-52 ste11:: NAT* | (Pitoniak et al. 2015) |
| PC4256 | *MAT***a** *ste4 FUS1-lacZ FUS1-HIS3 ura3-52 leu2::HYG rsr1::NAT* | (Pitoniak et al. 2015) |
| PC4805 | *MAT***a** *ste4 FUS1-lacZ FUS1-HIS3 ura3-52 BEM4$^{16-99}$-HA:: kanMX6* | (Pitoniak et al. 2015) |
| PC4807 | *MAT***a** *ste4 FUS1-lacZ FUS1-HIS3 ura3-52 BEM4$^{99-200}$-HA:: kanMX6* | (Pitoniak et al. 2015) |
| PC4809 | *MAT***a** *ste4 FUS1-lacZ FUS1-HIS3 ura3-52 BEM4$^{200-300}$-HA:: kanMX6* | (Pitoniak et al. 2015) |
| PC4811 | *MAT***a** *ste4 FUS1-lacZ FUS1-HIS3 ura3-52 BEM$^{300-400}$-HA:: kanMX6* | (Pitoniak et al. 2015) |
| PC5802 | *S288c* JMY1585 *MAT**a** rsp5::HIS3 pDsred415-rsp5ww2* | (Zhao et al. 2013b) |
| PC5803 | *S288c* JMY1585 *MAT**a** rsp5::HIS3 pnTAP416-rsp5ww3* | (Zhao et al. 2013b) |
| PC5851 | *MAT**a** ura3-52 his3-200 ade2-101 his3-200 leu2-1* | (Kono et al. 2012) |
| PC5852 | *MAT**a** ura3-52 his3-200 ade2-101 his3-200 leu2-1 cim3-1* | (Kono et al. 2012) |
| PC5024 | *MAT***a** *ura3-52 ste11::NAT* | (Pitoniak et al. 2015) |
| PC6017 | *MAT***a** *can1Δ::Ste2pr-spHIS5 lyp1Δ::Ste3pr-LEU2 his3::hisG* leu2Δ0 *ura3Δ0* | (Ryan et al. 2012) |
| PC6539 | *MAT***a** *ste4 FUS1-lacZ FUS1-HIS3 ura3-52 cdc42::NAT* pRS316-GFP-linker-CDC42 | This study |
| PC6591 | *MAT***a** *ura3-5 leu2* | This study |
| PC6810 | *MAT***a** *ura3-52 leu2 ssk1* | This study |
| PC6604 | *MAT***a** *ura3-52 leu2 ssk1 ste11::NAT* | This study |
| PC6680 | *MAT***a** *ura3-52 ssk1 bem1::NAT pRS316-BEM1* | (Basu et al. 2020) |
| PC7034 | *MAT***a** *ste4 FUS1-lacZ FUS1-HIS3 ura3-52 gic1:: kanMX6* | (Prabhakar et al. 2020) |
| PC7043 | *MAT***a** *ste4 FUS1-lacZ FUS1-HIS3 ura3-52 gic2:: NAT* | (Prabhakar et al. 2020) |
| PC7044 | *MAT***a** *ste4 FUS1-lacZ FUS1-HIS3 ura3-52 gic1:: kanMX6 gic2:: NAT* | (Prabhakar et al. 2020) |
| PC7086 | *MAT***a** *ste4 FUS1-lacZ FUS1-HIS3 ura3-52 bni1::NAT* | This study |
| PC7365 | MATa *ura3-52 CDC3-mCherry::HYG* | (Prabhakar et al. 2020) |
| PC7657 | *S288c MATα his3Δ0, leu2Δ0, met15Δ0, ura3Δ0 ydj1::kanMX6* | Research Genetics |

All strains are Σ1278b background unless otherwise indicated.

*Strains from an ordered deletion collection described in (Ryan et al. 2012) were also used.

**Table S2. Plasmids used in the study.**

| Name | Description | Reference |
|---|---|---|
| PC2207 | pRS316 | (Sikorski and Hieter 1989) |
| PC6455 | pRS306-GFP-linker-CDC42P (pDLB3609) | (Irazoqui et al. 2003) |
| PC6454 | pRS316-GFP-linker-CDC42 | (Basu et al. 2020) |
| PC7350 | pGFP-Cdc42$^{C188S}$ | This study |
| PC7349 | pRS316-GAL1-GFP-Cdc42p | This study |
| PC7455 | pGFP-Cdc42$^{D57Y}$ | This study |
| PC7458 | pGFP-Cdc42$^{Q61L}$ | This study |
| PC7504 | pGFP-Cdc42$^{K5R,K16R}$ | This study |
| PC7505 | pGFP-Cdc42$^{K5R,K16R,K123R,K128R,K166R}$ | This study |
| PC7506 | pGFP-Cdc42$^{K16R,K166R}$ | This study |
| PC7507 | pGFP-Cdc42$^{K5R,K16R.K94R,K96R}$ | This study |
| PC7508 | pGFP-Cdc42$^{K94R,K96R}$ | This study |
| PC7509 | pGFP-Cdc42$^{K5R,K16R,K94R,K96R,K123R,K128R,K166R}$ | This study |
| PC7511 | pGFP-Cdc42$^{K150R,K153R}$ | This study |
| PC7512 | pGFP-Cdc42$^{K5R,K16R,K150R,K153R,K166R}$ | This study |
| PC7513 | pGFP-Cdc42$^{K5R,K16R,K94R, K96R, K123R,K128R,K150R,K153R,K166R}$ | This study |
| PC7518 | pGFP-Cdc42$^{K166R}$ | This study |
| PC7520 | pGFP-Cdc42$^{K183R,K184R,K186R,K187R}$ | This study |
| PC7521 | pGFP-Cdc42$^{K5R,K16R,K94R, K96R,}$ K123R,K128R,K150R,K153R,K166R,K183R,K184R,K186R,K187R | This study |
| PC7522 | pGFP-linker-KKSKKCTIL | This study |
| PC7571 | p6xHis-linker-Cdc42 | This study |
| PC7572 | p6xHis-linker-Cdc42$^{K5R,K16R,K94R, K96R,}$ K123R,K128R,K150R,K153R,K166R,K183R,K184R,K186R,K187R | This study |
| PC7633 | pGFP-Cdc42$^{K5R,K94R, K96R,}$ K123R,K128R,K150R,K153R,K166R,K183R,K184R,K186R,K187R | This study |
| PC7635 | pGFP-Cdc42$^{K5R, K123R,K128R,K166R}$ | This study |
| PC7636 | pGFP-Cdc42$^{K5R, K94R,K96R}$ | This study |
| PC7637 | pGFP-Cdc42$^{K5R, K150R,K153R,K166R}$ | This study |
| PC7637 | pGFP-Cdc42$^{Q61L,K166R}$ | This study |
| PC7651 | pGFP-Cdc42$^{K5R, Q61L,K123R,K128R,K166R}$ | This study |
| PC7654 | pGFP-Cdc42$^{K5R, Q61L,K94R,K128R}$ | This study |
| PC7662 | pGFP-Cdc42$^{Q61L,K94R,K128R}$ | This study |
| PC7664 | pGFP-Cdc42$^{Q61L,K150R,K153R}$ | This study |
| PC7666 | pGFP-Cdc42 $^{K5R,K16R,Q61L,K94R, K96R,}$ K123R,K128R,K150R,K153R,K166R,K183R,K184R,K186R,K187R | This study |
| PC7667 | pGFP-$^{Cdc42K5R,Q61L,K150R,K153R,K166R}$ | This study |
| PC7633 | pGFP-Cdc42$^{K5R,K94R, K96R,}$ K123R,K128R,K150R,K153R,K166R,K183R,K184R,K186R,K187R | This study |
| PC7635 | pGFP-Cdc42$^{K5R, K123R,K128R,K166R}$ | This study |
| PC7636 | pGFP-Cdc42$^{K5R, K94R,K96R}$ | This study |
| PC7637 | pGFP-Cdc42$^{K5R, K150R,K153R,K166R}$ | This study |
| PC7637 | pGFP-Cdc42$^{Q61L,K166R}$ | This study |
| PC7651 | pGFP-Cdc42$^{K5R, Q61L,K123R,K128R,K166R}$ | This study |
| PC7654 | pGFP-Cdc42$^{K5R, Q61L,K94R,K128R}$ | This study |
| PC7662 | pGFP-Cdc42$^{Q61L,K94R,K128R}$ | This study |
| PC7664 | pGFP-Cdc42$^{Q61L,K150R,K153R}$ | This study |
| PC7666 | pGFP-Cdc42 $^{K5R,K16R,Q61L,K94R, K96R,}$ K123R,K128R,K150R,K153R,K166R,K183R,K184R,K186R,K187R | This study |
| PC7667 | pGFP-$^{Cdc42K5R,Q61L,K150R,K153R,K166R}$ | This study |

**Table S3. Primers used in the study**

| Name | Sequence (5´-3´) | Approach |
|---|---|---|
| C188S F1 | *GTTATCAAGAAAAGTAAAAAAAGTACAATTTTGTAGTCATATTAGTA* | SDM |
| C188S R1 | *TACTAATATGACTACAAAATTGTACTTTTTTTACTTTTCTTGATAAC* | SDM |
| D57Y F1 | *CATATACGTTAGGTTTGTTTTATACGGCCGGTCAAGAAGATTA* | SDM |
| D57Y R1 | *TAATCTTCTTGACCGGCCGTATAAAACAAACCTAACGTATATG* | SDM |
| G12V-F | *TGTGTTGTTGTCGGTGATGTTGCTGTTGGGAAAACGTGCCT* | SDM |
| G12V-R | *AGGCACGTTTTCCCAACAGCAACATCACCGACAACAACACA* | SDM |
| Q61L F1 | *GTTTGTTTGATACGGCCGGTCTAGAAGATTACGATCGATTGAG* | SDM |
| Q61L R1 | *CTCAATCGATCGTAATCTTCTAGACCGGCCGTATCAAACAAAC* | SDM |
| K166R F1 | *ACTAACACAACGCGGTTTGAGGAATGTATTCGATGAAGCT* | SDM |
| K166R R1 | *AGCTTCATCGAATACATTCCTCAAACCGCGTTGTGTTAGT* | SDM |
| K5R K16R F1 | *TGCGGCCGCGCACCACCACGACGACTAGTACACCCACAAACGCTAAGGTGTGTTGTTG TCGGTGATGGTGCTGTTGGGAGAACGTGCCTTCTAATCTCCTATA* | HR |
| K123R K128R R1 | *AACCTTGTTCTGATGTAATCGGACGTAATCTTTGTCTTTGCAACCTCTCGATGATTACCC TGTCATCCCTTAGATCAATCTGCG* | HR |
| K94R K96R R1 | *TGGTACACCTGGACAATGGTGATGTACTTCAGGGAACCATCTTTCTCTAACGTTTTCAAA AGAGGGTGGGGA* | HR |
| K150R K153R R1 | *CAAACCGCGTTGTGTTAGTGCCGAACACTCGACATATCTTACTGCTCTCAGTTCTCTTGC TAACCTGGA* | HR |
| K183R K184R K186R K187R F1 | *TTGGAGCCTCCTGTTATCAGGAGAAGTAGAAGATGTACAATTTTGTAGTC* | SDM |
| K183R K184R K186R K187R R1 | *GACTACAAAATTGTACATCTTCTACTTCTCCTGATAACAGGAGGCTCCAA* | SDM |
| GFP by His6x F1 | *AAATAAACGTATTAGGTCTTCCACAAAATGAGCGGCCGCATGCATCATCATCATCATCAT GGATCCTGCGGCCGCGCACCACCACGACGACTAGTACACCCA* | HR |
| GFP by His6x R1 | *AAATAAACGTATTAGGTCTTCCACAAAATGAGCGGCCGCATG* | HR |
| GFP-stop-F | *ATGGATGAACTATACAAGTAGTCCTGCGGCCGCGCACCA* | SDM |
| GFP-stop-R | *TGGTGCGCGGCCGCAGGACTACTTGTATAGTTCATCCAT* | SDM |
| GFP-linker-KKSKK-CTIL-F | *GGATCCTGCGGCCGCGCACCACCACGACGACTAGTACACCCAAAGAAAAGT AAAAAATGTACAATTTTGTAAAGAAAAGTAAAAAA* | HR |
| GFP-linker-KKSKK-CTIL-R | *ACTACAAAATTGCACATTTTTTACTTTTCTTTACAAAATTGTACATTTTTTACTTTTCTTTGG GTGTACTAGTCGTCGTGGTGGTGCGCGGCCGCAGGATCC* | HR |
| CDC42-F | *GTTGTTGTCGGTGATGGTGC* | qPCR |
| CDC42-R | *GTTGGAACATAGTCGGCTGGA* | qPCR |
| GFP-F | *CAATTGGCGATGGCCCTGTC3* | qPCR |
| GFP-R | *GCCATGTGTAATCCCAGCAGC* | qPCR |
| ACT1-F | *TGGATTCCGGTGATGGTGTT* | qPCR |
| ACT1-R | *CGGCCAAATCGATTCTCAA* | qPCR |

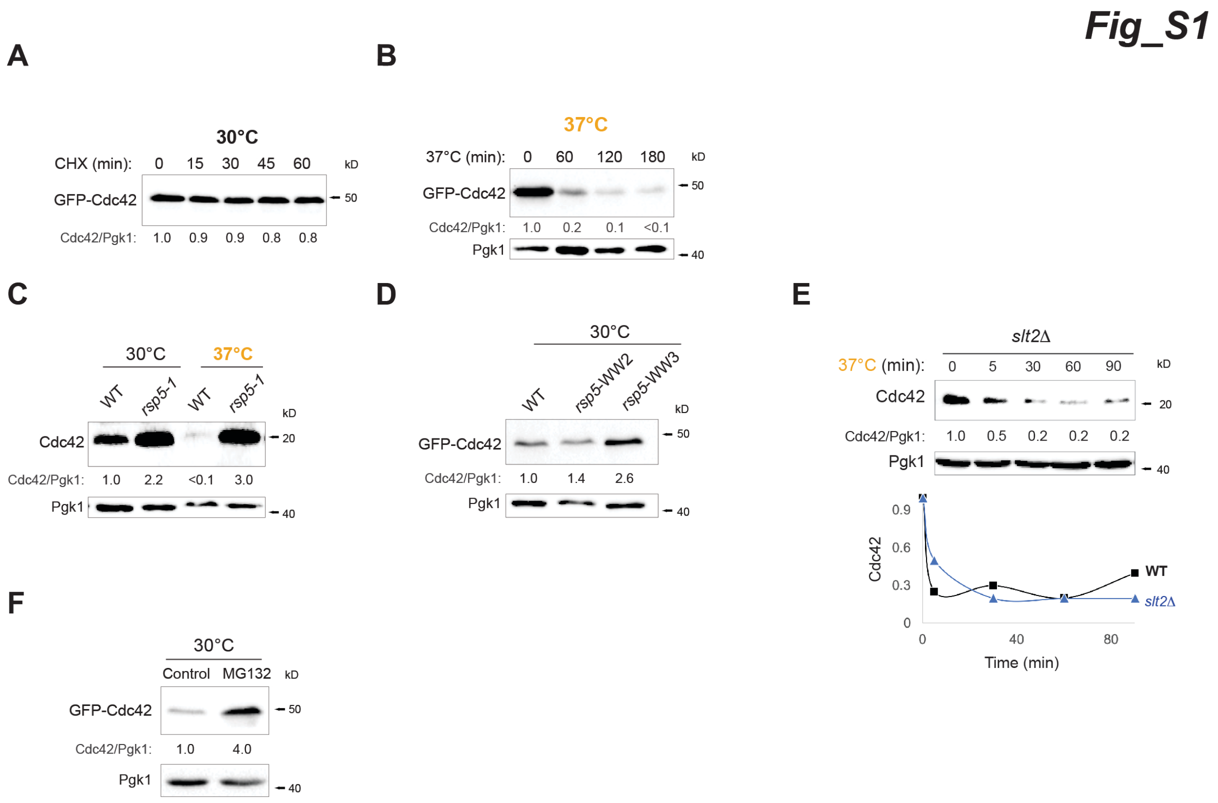


**Figure S1. The WW domains of Rsp5p and the proteasome are required for the degradation of Cdc42p. A)** WT cells were grown in media containing 25 µg/ml of CHX and analyzed by immunoblotting at the indicated time points. Pgk1p levels are the same as in Fig. 1B. Anti-GFP and anti-Pgk1p antibodies were used. Numbers indicate relative GFP-Cdc42p compared to Pgk1p. **B)** Cdc42p levels in WT cells were grown at 30°C for 4 h and shifted to 37°C. Samples were analyzed at the indicated time points. **C)** Endogenous Cdc42p levels of extracts prepared from WT and *rsp5-1* cells at 30°C and at 37°C. Anti-Cdc42p and anti-Pgk1p antibodies were used. Relative Cdc42p amount is indicated in numbers bellow the blot. **D)** GFP-Cdc42p levels were analyzed in wildtype and two alleles or Rsp5 which contains point mutations in two WW domains, *rsp5*-WW*2* (Rsp5$^{W359G}$) and *rsp5*- WW*3* (Rsp5$^{W451G}$). Anti-GFP and Pgk1p antibodies were used. Relative GFP-Cdc42p amount is indicated in numbers bellow the blot. **E)** WT and *slt2*Δ cells were grown at 30°C for 5 h and shifted to 37°C. Samples were analyzed at the indicated time points. Proteins extracts were detected using anti-Cdc42p and anti-Pgk1 antibodies. Graph represents relative levels of Cdc42p compared to Pgk1p. **F)** WT cells containing GFP-Cdc42p supplemented with 0.5% ethanol (control) or 75µM MG132 (MG132) were analyzed by immunoblotting. Anti-GFP and Pgk1p antibodies were used. Relative GFP-Cdc42p amount is indicated in numbers bellow the blot.

Fig_S2

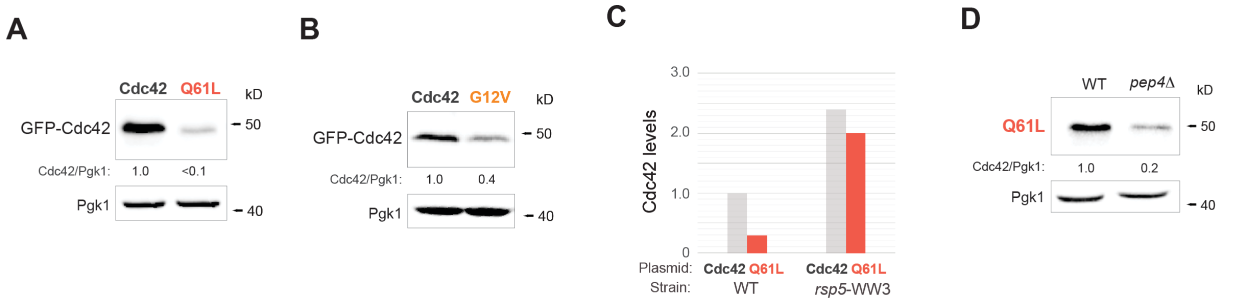


**Figure S2. GTP-locked versions of Cdc42p are found at low levels independent of the vacuolar protease Pep4p. A)** Levels of GFP-Cdc42p and GFP-Cdc42p$^{Q61L}$ in wild-type (WT, PC538) cells. Numbers indicate relative protein quantification. Anti-GFP and anti-Pgk1p antibodies were used. **B)** Levels of GFP-Cdc42p, GFP-Cdc42p$^{G12V}$ in WT (PC538) cells. Numbers indicate relative protein quantification. Anti-GFP and anti-Pgk1p antibodies were used. **C)** GFP-Cdc42p (Cdc42p) and GFP-Cdc42p$^{Q61L}$ (Q61L) relative levels in wild-type (WT) and in *rsp5-ww3* (Rsp5p$^{W451G}$) cells. **D)** Levels of GFP-Cdc42p$^{Q61L}$ in WT cells and cells lacking the vacuolar protease Pep4p. Numbers indicate relative Cdc42p quantification. Anti-GFP and anti-Pgk1p antibodies were used.

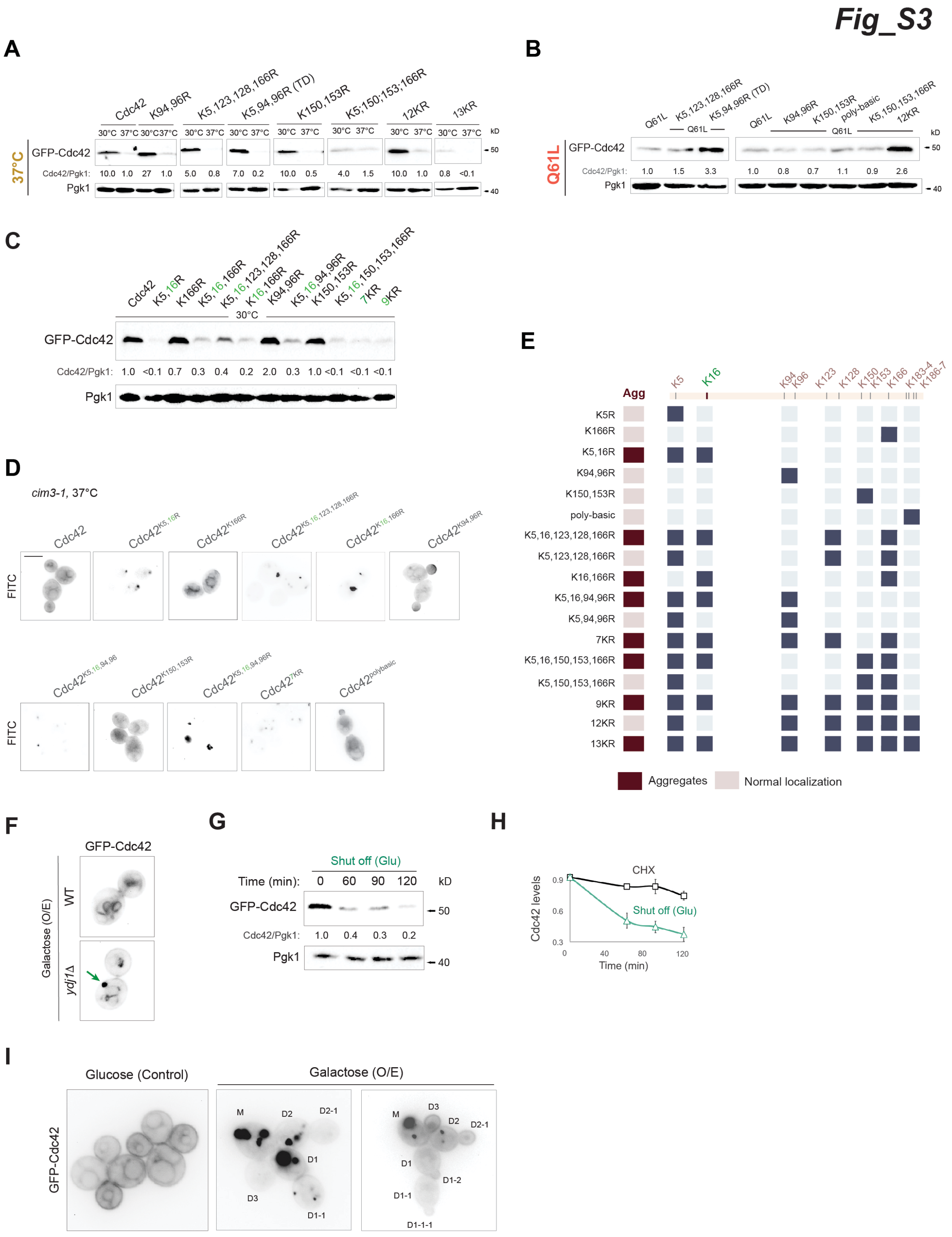
Fig_S3
A
Cdc42
K94,96R
K5,123,128,166R
K5,94,96R (TD)
K150,153R
K5;150;153;166R
12KR
13KR
37°C
GFP-Cdc42
Cdc42/Pgk1:
Pgk1
B
Q61L
GFP-Cdc42
Cdc42/Pgk1:
Pgk1
C
GFP-Cdc42
Cdc42/Pgk1:
Pgk1
30°C
D
cim3-1, 37°C
FITC
E
Agg
Aggregates
Normal localization
F
GFP-Cdc42
WT
ydj1Δ
Galactose (O/E)
G
Shut off (Glu)
Time (min):
GFP-Cdc42
Cdc42/Pgk1:
Pgk1
H
CHX
Shut off (Glu)
Cdc42 levels
Time (min)
I
Glucose (Control)
Galactose (O/E)
GFP-Cdc42

**Figure S3. Role of specific lysines in Cdc42p levels. K16R mutation and overexpression of *CDC42* promote the formation of aggregates in mother cells. A)** GFP-Cdc42p levels of the indicated alleles grown and at 30°C for 5 h (30°C) or shifted to 37°C for 1 h (37°C). Anti-GFP and anti-Pgk1p antibodies were used. Numbers indicate the relative levels of GFP-Cdc42p compared to Pgk1p. **B)** GFP-Cdc42p levels of wild-type (WT) cells expressing the indicated Cdc42p alleles grown at 30°C for 6 h (K5,94,96R,TD, turnover deficient). Anti-GFP and anti-Pgk1p antibodies were used. Numbers indicate the relative levels of GFP-Cdc42p compared to Pgk1p. **C)** Immunoblots of lysates of Cdc42p alleles grown at 30°C for 6 h. Proteins extracts were detected using anti-GFP and anti-Pgk1p antibodies. Numbers indicate relative levels of GFP-Cdc42p compared to Pgk1p. **D)** Fluorescence microscopy of *cim3-1* cells expressing the indicated alleles grown to mid-log phase at 30°C and shifted to 37°C for 1 h. **E)** Heat map indicates Cdc42p alleles which formed aggregates in *cim3-1* cells grown at 37°C for 1 h. **F)** WT cells and cells lacking Ydj1p expressing $pP_{GAL1}$-GFP-linker-*CDC42P* were grown in YEP-GAL for 3 h and analyzed by fluorescence microscopy. Scale bar, 5 µm. **G)** WT cells expressing $pP_{GAL1}$-GFP-linker-*CDC42P* were grown in YEP-GAL and shifted to glucose medium (YEPD), samples were collected and analyzed at the indicated time points. Anti-GFP and anti-Pgk1p antibodies were used. Numbers indicate relative GFP-Cdc42p levels compared to Pgk1p. **H)** Quantitative analysis of GFP-Cdc42p turnover in cells grown in CHX(CHX, grey, panel 2A), and cells expressing $pP_{GAL1}$-GFP-linker-*CDC42P* grown in YEP-GAL and shifted to YEPD (Shut off, green, panel S3G. Error bars represent biological duplicates. **I)** WT cells expressing $pP_{GAL1}$-GFP-linker-*CDC42P* were grown in YEPD (Glucose, control) or YEP-GAL (Galactose; O/E, overexpression) for 8 h and analyzed by fluorescence microscopy. M refers to mother cell, D to daughter cell. Scale bar, 5 µm

Fig_S4

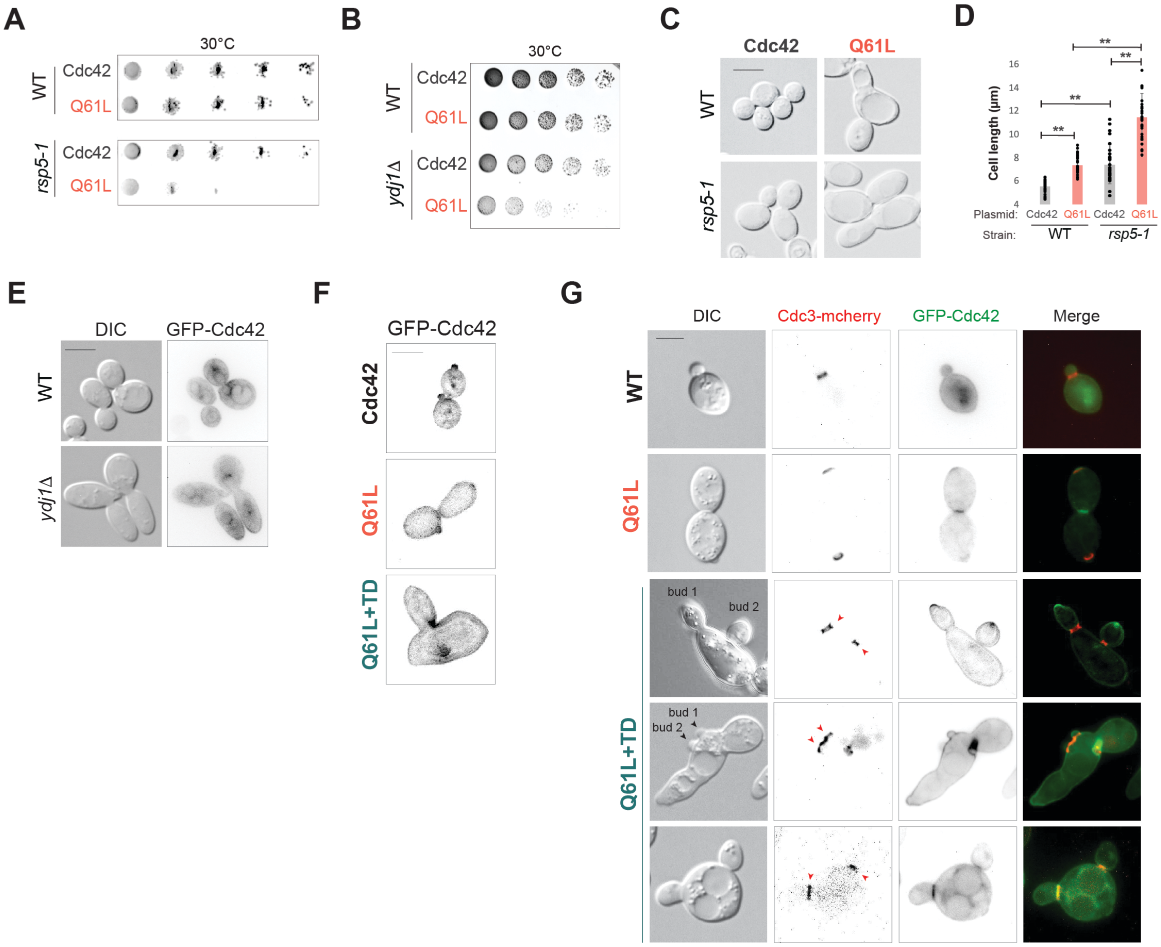


**Figure S4. Turnover-deficient version of Cdc42p (Cdc42$^{Q61L+TD}$) affects polarity and viability. A)** Serial dilutions of WT and *rsp5-1* cells expressing GFP-Cdc42p or GFP-Cdc42$^{Q61L}$. Cells were grown for 2 days at 30°C on SD-URA media. **B)** Serial dilutions of WT and *ydj1*Δ cells expressing GFP-Cdc42p or GFP-Cdc42$^{Q61L}$. Cells were grown for 2 days at 30°C on SD-URA media. **C)** Microscopic examination of WT and *rsp5-1* cells expressing GFP-Cdc42p or GFP-Cdc42p$^{Q61L}$. Scale bar, 5 µm. **D)** Quantification of cell length of cells from WT and *rsp5-1* cells expressing GFP-Cdc42p or GFP-Cdc42$^{Q61L}$. N >25, asterisk P-value <0.01. **E)** Fluorescence microscopy of WT and *ydj1*Δ cells expressing GFP-Cdc42p. Scale bar, 5 µm. **F)** Microscopic examination of cells expressing GFP-Cdc42p (Cdc42), GFP-Cdc42p$^{Q61L}$ (Q61L) or GFP-Cdc42p$^{Q61L+K5,94,96R}$ (Q61L+TD). Scale bar, 5 µm. **G)** Fluorescence microscopy of Cdc3-mcherry cells expressing GFP-Cdc42p (Cdc42), GFP-Cdc42p$^{Q61L}$ (Q61L) or GFP-Cdc42p$^{Q61L+K5,94,96R}$ (Q61L+TD). Cdc3-mcherry (red) is a septin ring marker. Scale bar, 5 µm.

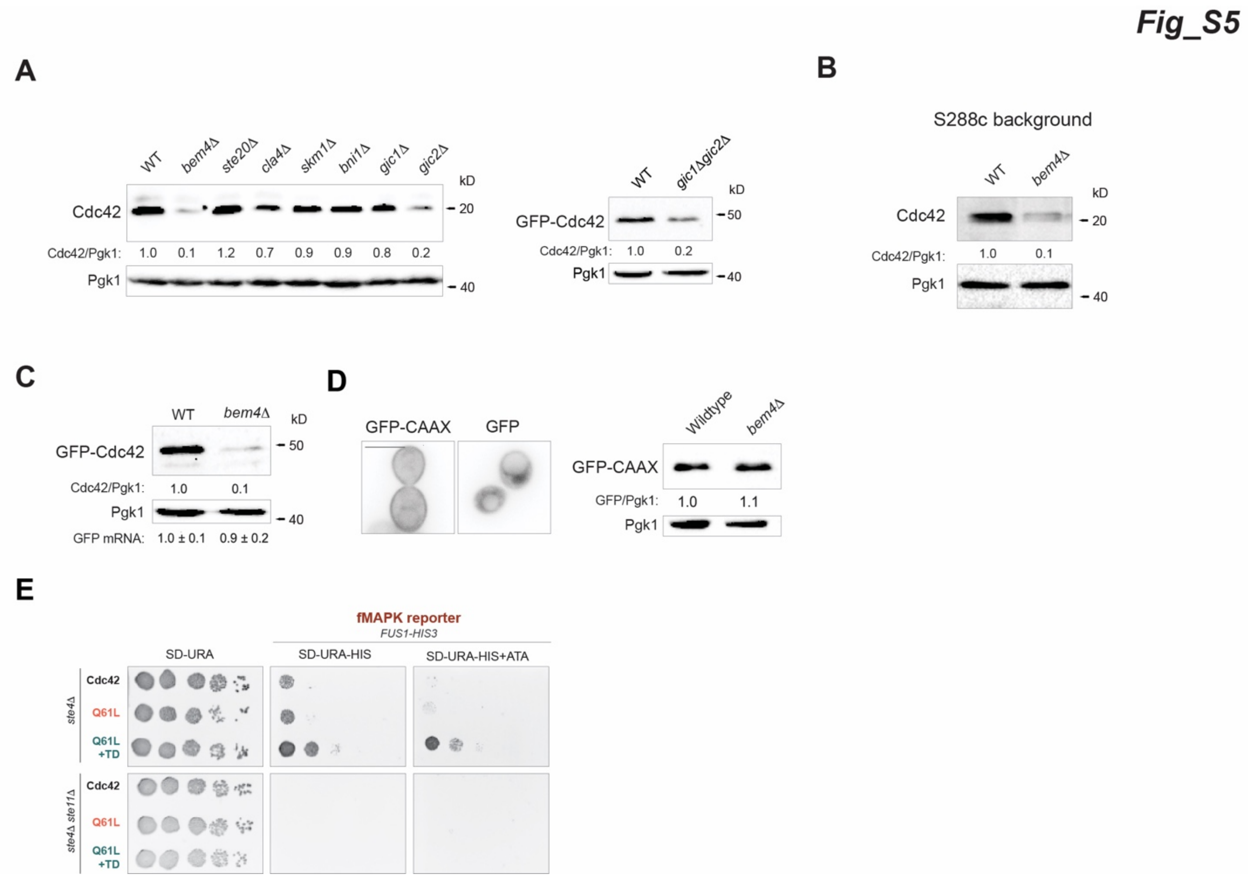


**Figure S5. Cells lacking the fMAPK scaffold Bem4p show reduced levels of Cdc42p. A)** Cdc42p or GFP-Cdc42p levels of extracts prepared from wild-type (WT, PC538) cells or interacting-proteins mutants. Anti-Cdc42p, anti-GFP and anti-Pgk1 antibodies were used. Numbers indicate relative levels Cdc42p or GFP-Cdc42p compared to Pgk1p. **B)** WT cells from the S288c background were analyzed as in panel A**. C)** WT cells and *bem4*Δ mutant expressing GFP-Cdc42p analyzed by immunoblotting. Anti-GFP and anti-Pgk1p antibodies were used. Cdc42/Pgk1 ratio indicates GFP-Cdc42p levels normalized to Pgk1. *GFP mRNA* levels were detected by RT-PCR. Standard deviation was generated by analysis of two independent trials. **D)** Left panel, fluorescent examination of cells carrying a pGFP-KKSKKCTIL or pGFP plasmids. Scale bar, 5 µm. Right panel, GFP-KKSKKCTIL protein levels in wild-type cells and in the *bem4*Δ mutant. Anti-GFP and anti-Pgk1p antibodies were used. Numbers indicate relative amount of GFP-KKSKKCTIL compared to Pgk1p. **E)** Serial dilutions of the *ste4*Δ mutant containing the growth reporter *FUS1-HIS3* and expressing the indicated alleles of Cdc42p were growth on SD-URA plates, SD-URA-HIS, and SD-URA-HIS + 1.6µM ATA (fMAPK reporter).
